# MycorrhizaFinder: a user-friendly machine learning tool to quantify endomycorrhizal colonisation of real-world roots

**DOI:** 10.64898/2026.03.04.709422

**Authors:** Jill Kowal, Robin Upham, Shaun Buckley, Adam Kiani, Morgan Rickards, Ella Serpell, Martin I. Bidartondo, Edouard Evangelisti, Sebastian Schornack, James Sibbit, Kevin P. Treder, Simon Weidinger, Laura M. Suz

## Abstract

**Background and aims:** Root colonisation by endomycorrhizal fungi has long been one of the most widely used metrics in mycorrhizal studies. However, due to the significant time required to assess colonisation using traditional microscope techniques, studies of colonisation at large scales are impractical. AI-powered approaches may increase output and facilitate ecosystem assessments.

**Methods:** We trained an AI-powered tool (MycorrhizaFinder) on field roots from diverse grasslands and heathlands hosting common Northern European plants with a range of arbuscular (AM) and ericoid mycorrhizal (ErM) fungal structures, and dark septate endophytes (DSE), also common in field-sourced roots. We incorporated a user-customized confidence threshold to encourage the user to engage with inevitable morphological ambiguities, in e.g. ErM and DSE. A Macro F_1_ statistic was used to assess the tool’s development.

**Results:** We provide a sample workflow from root processing and microscope slide scanning to semi-automated model training and performance evaluation. Without human supervision, our automated baseline Macro F_1_ is 66% for arbuscular and at 57% for ericoid mycorrhizal colonisation assessment.

**Conclusion:** MycorrhizaFinder is user friendly, requires no programming skills and offers flexibility for advanced agronomists or ecologists who wish to train the tool using their own labelled mycorrhizal root datasets, including images acquired from different instruments or staining protocols. This adaptability allows users to customize the model for specific ecosystems or experimental designs. Leveraged with molecular identification and/or functional assessment of fungi, MycorrhizaFinder could support scalable and repeatable monitoring across ecosystems to assess mycorrhizal status and track land-use changes over time.

## Introduction

Most land plants form endomycorrhizas, defined by nutrient exchange between a plant root (‘host’) and beneficial fungi (Brundrett and Tedersoo, 2018). Roughly 85% of plants are colonised by intracellular hyphae of diverse Glomeromycotina and/or Mucoromycotina (Endogonales) fungi, forming arbuscular mycorrhizas (AM) or AM-like symbioses, with each subphylum having distinct morphological and nutritional functional traits (Hoysted et al. 2018, 2019). Ericaceae plants found in Northern European heathlands and also abundant as understory in woodlands are instead colonised by ericoid mycorrhizal (ErM) fungi. The Ericaceae constitute ∼1.5% of terrestrial hosts for members of the fungal phyla Ascomycota and Basidiomycota. Many ErM types have been described since the classical intercellular ascomycete coils were described by Read (1996); Kiheri et al. (2020) describe four, distinguishing them from DSE which have more heavily melanised septate hyphae. There is a further non-mycorrhizal fungal group, the dark septate endophytes (DSE), which is predominantly composed of Ascomycota. The ecological functions of DSE remain enigmatic (de Oliveira and Pereira 2025), but they are globally widespread and may be central to understanding some ecological systems (Netherway et al. 2024). A continuum of structures has also been observed between ErM and DSE in e.g. *Rhododendron* spp., drawing attention to both the morphological ambiguities that may arise when categorizing mycorrhizal type based on morphology alone, and unresolved biological functions of both groups (Vohník and Albrechtová 2011).

The extent of mycorrhizal colonisation is an informative, context-dependent indicator of functional outcomes for host plants and is responsive to edaphic changes and host secondary metabolites (Soudzilovskaia et al. 2015; Frew 2025). Direct manipulation experiments using colonisation as a response variable demonstrate mixed outcomes. Positive relationships between AM and ErM fungi colonisation levels with aboveground growth and resilience in field and laboratory experiments have been reported (Diaz et al. 2006; Kowal et al. 2016; Lethielleux-Juge 2025, Frew et al. 2025). A habitat scale study showed that AMF root colonisation and soil neutral lipid fatty acids (NFLA) were strongly related, raising the possibility of using colonisation as a proxy for soil AMF biomass (Barcelo et al. 2020). A Google Scholar search in May 2026 using the phrases ‘arbuscular mycorrhizal fungi colonisation assessments’ or ‘ericoid mycorrhizal fungi colonisation assessments’ yielded 24,300 and 5,000 articles since 2020, respectively, showing that colonisation constitutes a parameter widely used by researchers. Typically, colonisation is evaluated using microscopy, either by counting a number of grid intersections with colonised root fragments (5-10mm) using a microscope eyepiece reticle, or by estimating the proportion of root cells that are colonised (McGonigle et al. 1990, Kowal et al. 2020). While these approaches offer reasonable objectivity and reproducibility, they are labour-intensive and therefore impractical for ecosystem-scale assessments, which often require the evaluation of hundreds to thousands of roots samples.

As part of the Department for Environment, Food and Rural Affairs’ (Defra) England-wide Natural Capital and Ecosystem Assessment programme, we have been evaluating field-sourced roots for a nationwide baseline assessment of mycorrhizas across terrestrial habitats, including arable and horticultural land, grasslands, moorlands, woodlands, heathlands and bogs. To expedite the analysis of thousands of roots, we initially tested the suitability of AMFinder (Evangelisti et al. 2021), a tile-based computer vision tool using a shallow convolution neural network (CNN) architecture developed to quantify AM in four species of laboratory grown model-plant seedlings (*Nicotiana benthamiana, Medicago truncatula, Lotus japonicus* and *Oryza sativa*). The mycorrhizal fungal symbionts introduced to colonise these roots were from monoxenic cultures of four AM fungal genera - *Rhizophagus, Claroideoglomus, Rhizoglomus* and *Funneliformis*. The authors used three categories or classes to distinguish tiles (colonised, uncolonised and non-root) and predict mycorrhizal colonisation. AMFinder produced highly accurate predictions with high recall and precision (see Glossary) using these seedling roots. However, when tested with our high-resolution images of field-sourced root sections cleared and stained with similar ink-vinegar protocols, it often could not recognise roots, and only performed well in rare cases where hyphae were well-defined and roots were dominated by a single Glomeromycotina or Mucoromycotina (Endogonales) taxon and morphotype. Field roots usually present a complex mosaic of AM or ErM fungal types depending on host and fungal life stage, often with irregular hyphal diameters, and co-occurring with DSE and other non-mycorrhizal fungi, microsclerotia and bacteria. More advanced image analysis techniques developed to analyse AM colonisation, including deep learning or computer vision models, have so far only been trained on a limited set of plants such as crops (e.g. sorghum or soybeans) grown and inoculated in controlled settings (Kokkoris et al. 2019, Muta et al. 2022; Sciascia et al. 2023; Zhang et al. 2024).

Overall, despite the increasing availability of computer vision tools for rapid analysis of mycorrhizal colonisation, existing solutions rely on root images from controlled glasshouse or laboratory experiments with limited variation in host plant, fungal symbiont or level of disturbance of roots. This creates a significant gap in efforts to accelerate colonisation assessment of field-collected roots, which are far more complex than binary plant-fungus samples, exhibiting diverse ecological backgrounds, variable staining quality and a mixture of fungal structures. To address this gap, we developed MycorrhizaFinder (MF, available at https://doi.org/10.5281/zenodo.17979413), a computer-vision tool to assess the colonisation of field roots across plant species and endomycorrhizal types, able to differentiate common artifacts and disturbance arising from field collection (e.g. root image obstruction by clay particle debris), and for use across terrestrial ecosystems. Our main aim was to investigate and test appropriate model architectures which would allow us to: 1) add additional classes in the classification of AM fungi; 2) develop an alternative class model infrastructure focusing on ErM fungal colonisation that could be toggled within the same, unified user interface; 3) incorporate contextual information into the classification of each tile using the surrounding tiles; and 4) upgrade the user interface with all modelling, labelling and training streamlined in a single workflow with human supervision. Most significantly, MF aims to be accessible by researchers with no coding skills and for laboratory and field-sourced roots alike, paving the way for large-scale assessments. We do not intend to simplify the rich and diverse fungal morphological and ontogenetic still-life images shared in the accompanying dataset by distilling them into a handful of categories. Rather we present a semi-automated machine learning framework by which the researcher, e.g. biologist, ecologist or agronomist, can with high confidence settings, filter-out most uncolonised root cells and typical AM and ErM forms from scrutiny, allowing more time for the more complex and ambiguous fungal structures.

## Glossary

<u>Accuracy</u>: ratio of all correct predictions (true positives + true negatives) to the total number of samples. Performance models using accuracy may refer to true positives as ‘sensitivity’ and true negatives as ‘1 - specificity’.

Contextualisation: the ability of the tool to assess and incorporate the content of neighbouring regions of an image (tiles) when classifying one region.

Convolutional neural network (CNN): machine learning architecture that is especially well-suited to extracting information from patterns in images.

<u>Class</u>: one of the predefined categories a CNN can assign to an input. The final layer of the CNN produces a score or probability for each class at the tile level, and the prediction outcome is the class with the highest score.

<u>EfficientNet architecture</u>: A family of CNNs developed specifically for making use of image resources for computer vision tasks. This process is efficient (hence the name) at capturing strong relationships with a wide receptive field at a comparatively lower number of parameters to other state-of-the-art architectures, using a compound scaling coefficient to balance network depth, width and resolution. The EfficientNet model is generally larger and more complex than the CNN1 model, providing more learning capacity for the number of parameters it uses to capture rich global features and local fine-grained features. This is especially useful for images at a size of 126×126 or 252×252 pixels, where both capturing small details and an overall understanding of the image whilst utilizing the original image resolution are important for accurate classification. EfficientNetB5 is a medium-sized member of the family of models, with around 30 million trainable parameters.

<u>Precision</u>: a model performance metric that represents the impact of false positives (FP), calculated at a class level as the ratio of true positive (TP) identifications of said class, to all positive identifications of the class. i.e. it quantifies the prospect of a sample actually being positive when the model classifies it as positive.

<u>Recall</u>: a model performance metric that represents the impact of false negatives, calculated at a class level as the ratio of true positive identifications of said class to all tiles truly of that class. i.e. it quantifies the model’s ability to spot all positive instances.

<u>F_1_ Score</u>: a model performance metric that evaluates the performance of a particular class by balancing precision and recall. It is defined as the harmonic mean of precision and recall (Equation 1).

<u>Macro F_1_</u>: a model performance metric that combines the *F_1_* Scores for all classes, by taking their unweighted average, to provide insight into overall model performance (Equation 2). The <u>3-class Macro F_1_</u> refers to the three classes in AMFinder: *Colonised root section*, *Non-colonised root section,* and *Background*.

Max-vote system: the most probable class in the target tile is considered alongside that of four partially overlapping tiles and the target tile is assigned the class that is dominant among all five tiles.

React: a library for creating user interfaces for web applications, using the JavaScript programming language (Meta Open Source 2025).

## Materials and Methods

### Roots and Images

The roots used for training the model were obtained from field collected soil cores (15 cm deep, 2 cm wide) from agricultural fields, grasslands, heathlands and bogs across England, and *Calluna vulgaris* and *Erica tetralix* roots from sites in Norway. Soil samples were placed in a metal sieve (0.5 mm aperture size) and rinsed in tap water. Non-woody fresh roots were selected to yield a diverse range of root architectures, from straight to fine with complex branching (Fig. 1). All roots from putative AM hosts were stained in blue-ink and mounted on slides by modifying Kowal et al. (2020) (see Methods S1). Trypan blue solution (0.05% v/v) was used for ErM-host roots as we hypothesized that the ink may be less effective at adhering to intraradical hyphae. Root sections for both putative-AM and ErM hosts were 1cm long with variable surface according to root architectures. The slides were scanned to produce high resolution images using an Epredia 3DHISTECH Pannoramic MIDI II Rx brightfield digital slide scanner at 37x magnification (0.27 μm/pixel), but the tool can also adapt to jpg images acquired using a digital microscope (Evangelisti et al. 2021). To capture the three-dimensional, complex fungal network within the root cortex, an extended focus imaging approach was used to condense multiple-focal planes. Seven focus levels were used, where one step was around 0.2 μm. In total, root sections measuring approximately 1 cm in length from 133 unique roots (45 AM colonised, 88 ErM colonised) were selected for training after evaluating image quality and maximising quantity of tiles across classes (Table 1; complete training image library available at https://doi.org/10.5281/zenodo.17476686). Depending on the root, there were hundreds or thousands of tiles per root tissue area, i.e. non-background tiles, with each tile containing 252×252 pixels for AM and 126×126 for ErM tiles, set to align with typical root cell sizes.

**Figure 1.**
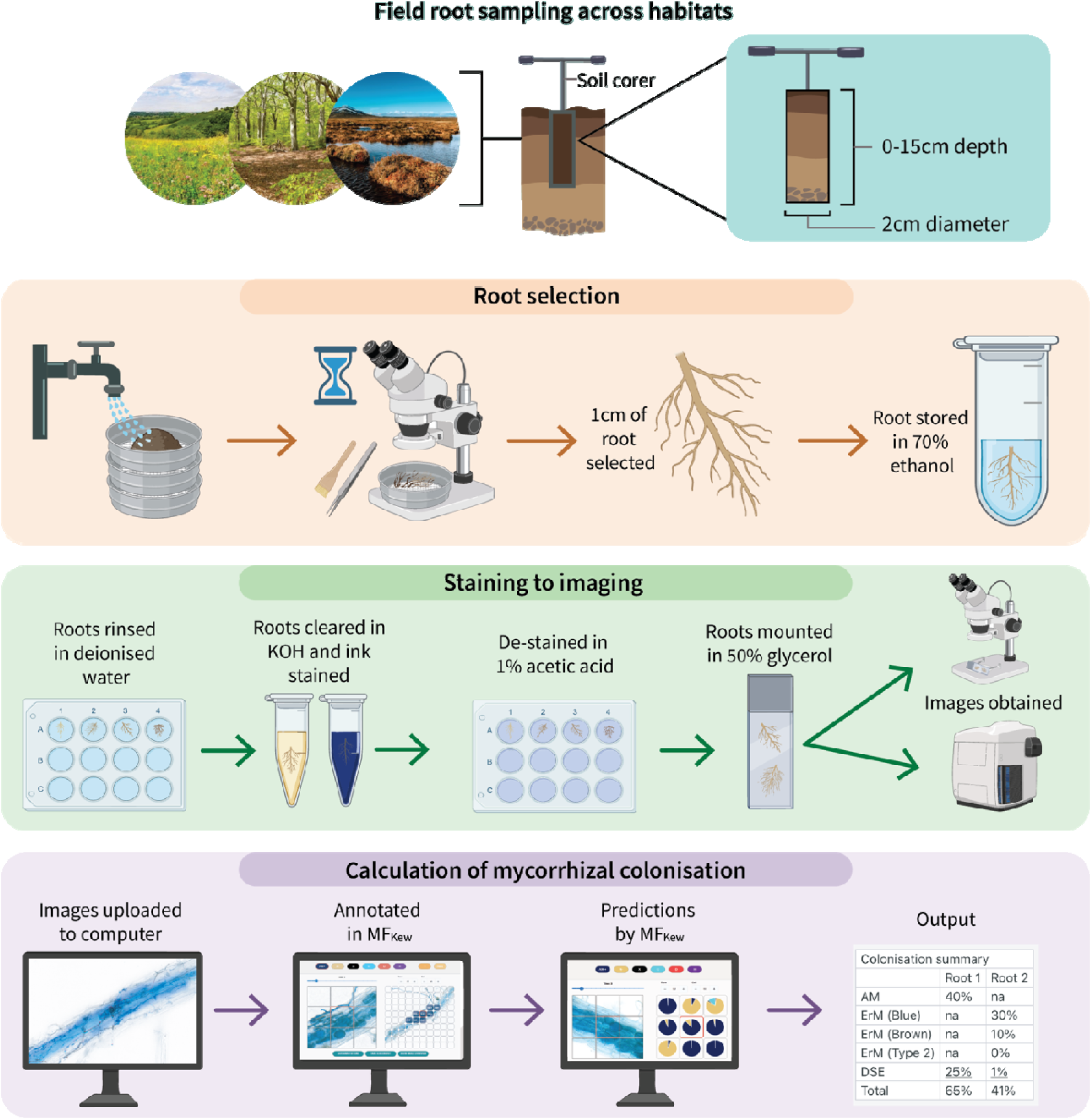
Workflow diagram showing field sampling, root selection, staining, imaging, and calculation of mycorrhizal colonisation. Bottom panel shows a simplified workflow of MycorrhizaFinder, from left to right: Image upload; Annotation of root section with nine tiles (training or editing); Predictions of tile class (pie-charts) with target tile outlined in red; Output, root level colonisation summary. Roots’ colonisation predictions can be processed in batches. Refer to https://doi.org/10.5281/zenodo.17979413 for further details.

**Table 1.** Summary of tile data classes used for training. Note: since a single root can contribute tiles from more than one class, the total number of roots used is less than the sum of the number of roots used for each class. AM: arbuscular mycorrhizal; DSE: dark septate endophyte.

| Model | Tile class / Abbreviation | Number of roots | Number of tiles | Percent of model tiles |
| --- | --- | --- | --- | --- |
| Arbuscular mycorrhiza | AM Colonised / AM<sup>+</sup> | 45 | 3,604 | 4.2 |
|  | DSE Colonised / DSE | 34 | 808 | 0.9 |
|  | AM+DSE Colonised / H | 24 | 276 | 0.3 |
|  | Uncolonised/ N- | 45 | 15,239 | 17.9 |
|  | Background / X | 46 | 63,736 | 74.7 |
|  | Unreadable / U | 37 | 1,705 | 2.0 |
| Ericoid mycorrhiza | Blue Coils Colonised / Bl+ | 76 | 2,344 | 2.8 |
|  | Brown Coils Colonised / Br+ | 76 | 2,485 | 3.0 |
|  | Type Two Colonised / T+ | 9 | 319 | 0.4 |
|  | DSE Colonised / DSE | 43 | 1,916 | 2.3 |
|  | Mixed ErM Colonised / HE+ | 56 | 314 | 0.4 |
|  | ErM+DSE Colonised / HD+ | 9 | 37 | < 0.1 |
|  | Uncolonised/ N- | 85 | 6,492 | 7.9 |
|  | Main Root /M | 85 | 12,838 | 15.6 |
|  | Background / X | 88 | 50,564 | 61.3 |
|  | Unreadable / U | 80 | 5,242 | 6.4 |

We also evaluated the model on a subset of images used to train AMFinder (but not MF) to test how the platform performed with images captured using a digital microscope more typical of plant science laboratories. We selected 19 root images compatible with the tile size parameters.*Distinction and quantification of mycorrhizal structures*

To accurately quantify mycorrhizal colonisation in field roots, we created more classes than the original three-class AMFinder (Evangelisti et al. 2021). This allowed our machine learning model to recognise a wider range of real-world root and fungal biological features than in the original training images used to build AMFinder. Real-world field roots present variable hyphal diameters and textures within diverse root architectures, co-occurrence of DSE, and cell structural noise from soil extraction. We modified the two original CNNs (CNN1 and CNN2) to expand the CNN1’s output layer from three classes in AMFinder to six in our model. Moreover, the CNN2’s functionality was incorporated into CNN1 for the ErM mode path, allowing it to differentiate between hyphal structures, described in Table 2. Images were augmented and datasets’ classes balanced using several standard machine learning techniques (see Methods S2).

**Table 2.**
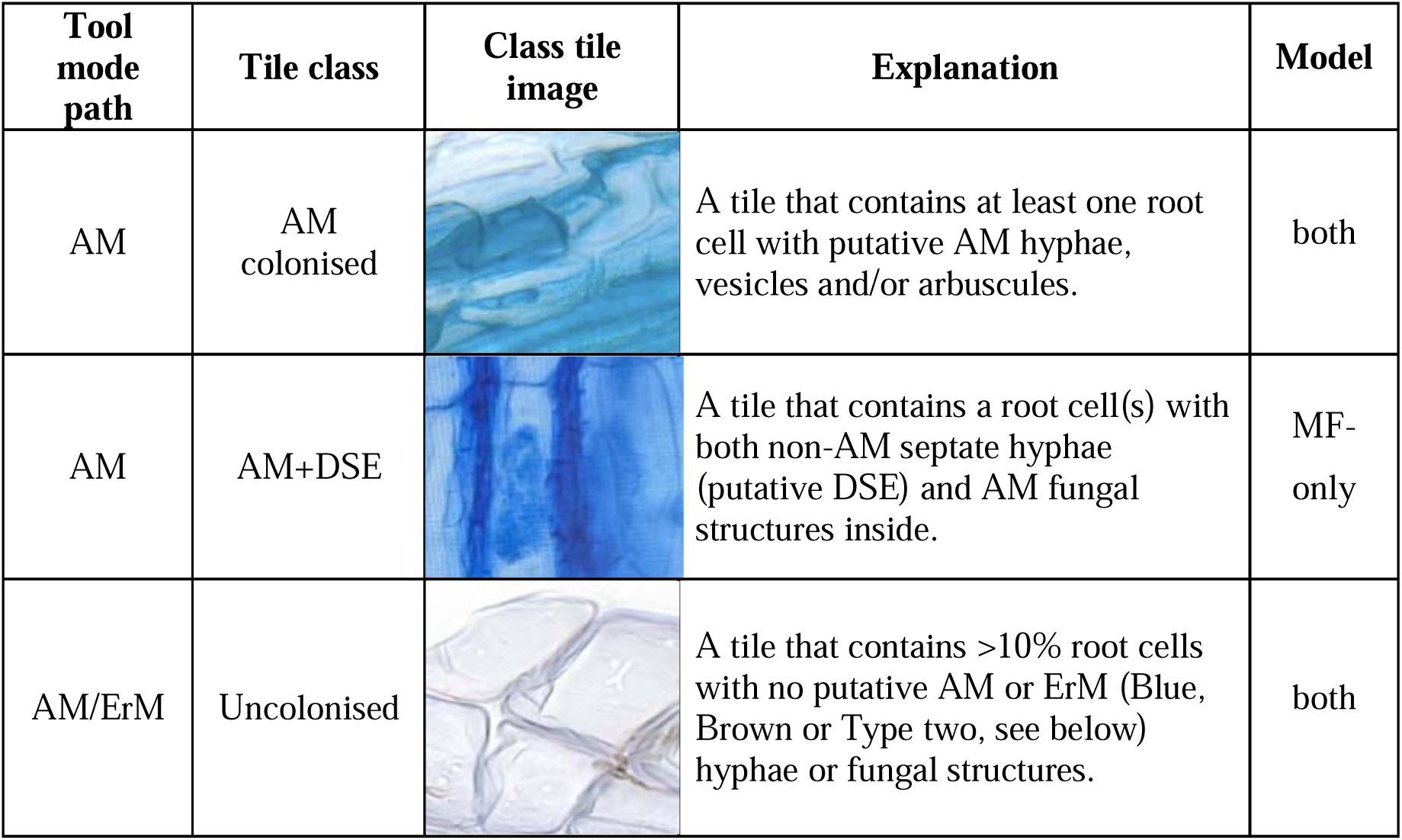
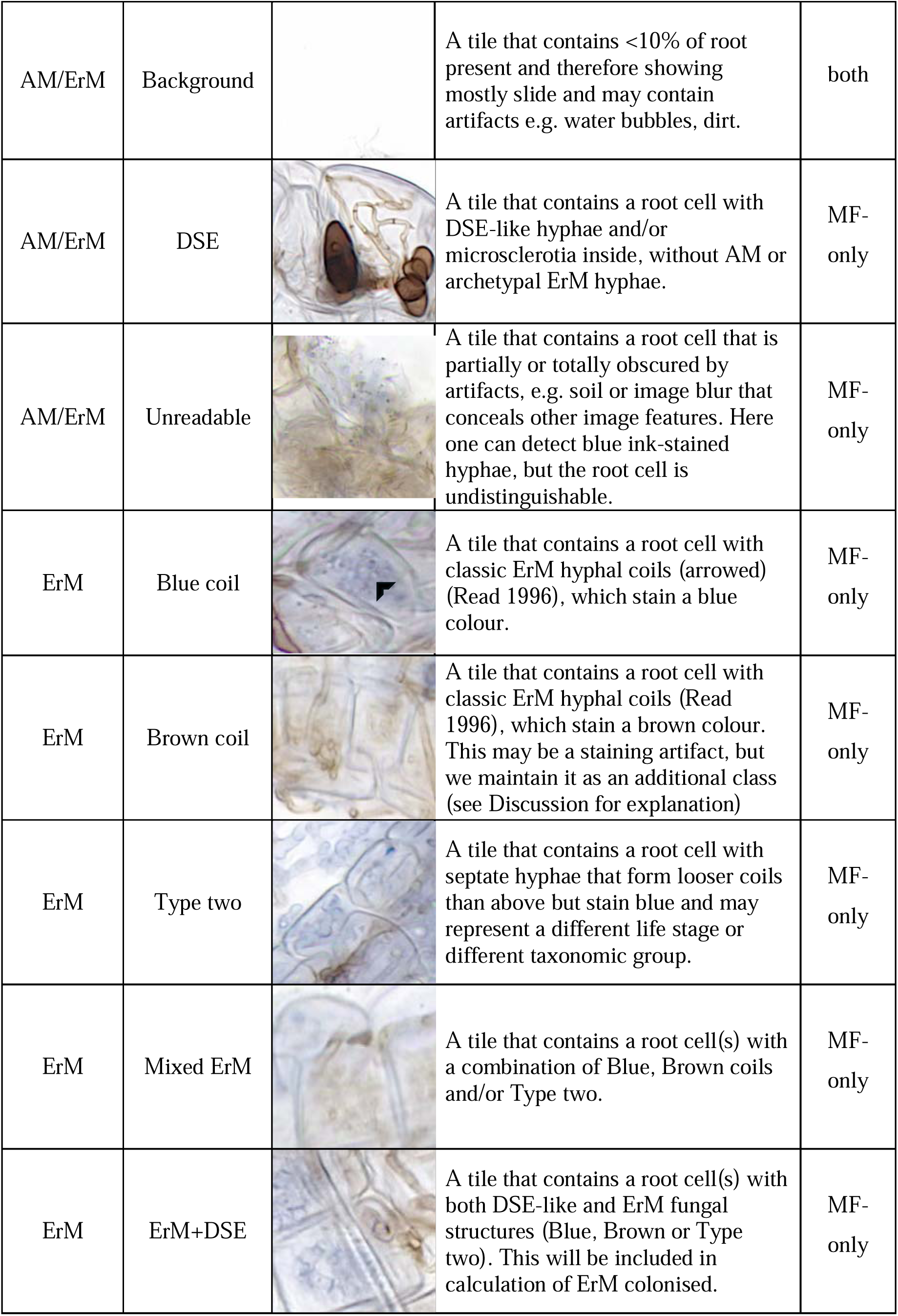
Tile classes utilised in the model output layer of AMFinder (Evangelisti et al. 2021) and MycorrhizaFinder (MF). Either ‘both’, AMFinder and the expanded MF models, or ‘MF-only’ for new features of arbuscular mycorrhiza (AM) and ericoid mycorrhiza (ErM) detection in roots. Class tile images show examples of defined class feature(s).

| Tool mode path | Tile class | Class tile image | Explanation | Model |
| --- | --- | --- | --- | --- |
| AM | AM colonised |  | A tile that contains at least one root cell with putative AM hyphae, vesicles and/or arbuscules. | both |
| AM | AM+DSE |  | A tile that contains a root cell(s) with both non-AM septate hyphae (putative DSE) and AM fungal structures inside. | MF-only |
| AM/ErM | Uncolonised |  | A tile that contains >10% root cells with no putative AM or ErM (Blue, Brown or Type two, see below) hyphae or fungal structures. | both |
| AM/ErM | Background |  | A tile that contains <10% of root present and therefore showing mostly slide and may contain artifacts e.g. water bubbles, dirt. | both |
| AM/ErM | DSE |  | A tile that contains a root cell with DSE-like hyphae and/or microsclerotia inside, without AM or archetypal ErM hyphae. | MF-only |
| AM/ErM | Unreadable |  | A tile that contains a root cell that is partially or totally obscured by artifacts, e.g. soil or image blur that conceals other image features. Here one can detect blue ink-stained hyphae, but the root cell is undistinguishable. | MF-only |
| ErM | Blue coil |  | A tile that contains a root cell with classic ErM hyphal coils (arrowed) (Read 1996), which stain a blue colour. | MF-only |
| ErM | Brown coil |  | A tile that contains a root cell with classic ErM hyphal coils (Read 1996), which stain a brown colour. This may be a staining artifact, but we maintain it as an additional class (see Discussion for explanation) | MF-only |
| ErM | Type two |  | A tile that contains a root cell with septate hyphae that form looser coils than above but stain blue and may represent a different life stage or different taxonomic group. | MF-only |
| ErM | Mixed ErM |  | A tile that contains a root cell(s) with a combination of Blue, Brown coils and/or Type two. | MF-only |
| ErM | ErM+DSE |  | A tile that contains a root cell(s) with both DSE-like and ErM fungal structures (Blue, Brown or Type two). This will be included in calculation of ErM colonised. | MF-only |

To accommodate a greater number of classes, we evaluated four model architectures - ResNet, ResNeXt, the AMFinder ‘CNN1’ model and EfficientNetB5 (Tan & Le 2019) - and the latter was ultimately selected for further analysis (see Methods S2). Model training and testing were also conducted using the original CNN1 architecture to establish a benchmark for comparative performance evaluation.

The dataset breakdown of tiles by class is presented in Table 1. There may be more than one root cell within a tile or a partial cell, as long as the cell walls are distinguishable and cover 10% of the tile (see MycorrhizaFinder Full User Documentation, https://doi.org/10.5281/zenodo.17979413).

A new class structure for Ericaceae plant family roots was added. We differentiated three ErM colonisation types based on morphology (*Blue coils*, *Brown coils* and *Type two*) (Table 2), for potential future analysis exploring biological and functional differences, recognising that these do not capture all the possibilities including a ErM-DSE continuum and hyphae related to ‘sheathed’ ErM host structures (Vohnik et al. 2012), which appear similar to Type two, but cannot be confirmed with cytological observations alone, i.e., without the whole-root context. We also distinguished ‘Main Root’ as the colonisation calculation is based only on young non-lignified roots.

### Quantification of mycorrhizal colonisation

We programmed the model to predict root colonisation in two steps. First, the model processes one tile at a time and computes the probability that the tile belongs to each class. Then, the contextual mode (default) incorporates four neighbouring tiles (with a diagonal overlap extending 75% from the central tile, Fig. S1) and uses a max-vote system to assign the class to each tile by combining majority predictions for all four neighbouring tiles with the target tile, thus generating a (new) class label for the central tile. As tiles and plant root cells are rarely aligned, this captured broader spatial context across neighbouring regions, mimicking the classification behaviour of a human when contextualizing observations, in this case to enable recognition of fungal structures within a complete root cell. Further details of the context feature are in Methods S2.

We included a feature allowing the user to switch on a threshold of confidence (the probability of the most probable class) above which, this context feature is disabled on a per-tile basis to save computing time. It defaults to a value of 1 as there is a systematic increase in overall predictive performance with the contextual feature enabled.

Once predictions are generated, they are required to be converted to annotations (definite class labels) to calculate the colonisation metrics. At this stage it is again possible to supply a threshold value of confidence, above which predictions are automatically converted to annotations of the most probable class and below which predictions are ignored. To solicit user recognition of the ambiguities inherent in these groupings, we suggest the user start at 80% (see user guide).

The calculation of colonisation is explained in Table 3. Each colonisation metric was defined as the ratio of tiles designated as belonging to a class(es), relative to the total number of tiles belonging to all root classes, excluding tiles which were background or unreadable.

**Table 3.** Definitions of colonisation metrics calculated by MycorrhizaFinder. The name of each class (*AMColonised*, (*AM+DSE*), etc.) refers to the number of tiles in the image belonging to the relevant class. For arbuscular mycorrhiza (AM), *TotalRoot* is defined as *Uncolonised* + *AMColonised* + *DSE* + (*AM+DSE)*; for ericoid mycorrhiza (ErM), it is defined as *Uncolonised* + *BlueCoils* + *BrownCoils* + *TypeTwo* + *DSE* + *MixedErM +* (*ErM+DSE)*.

| Colonisation type | Colonisation metric | Metric equation |
| --- | --- | --- |
| Arbuscular mycorrhiza | Arbuscular mycorrhizal | $\frac{AMColonised + (AM + DSE)}{TotalRoot}$ |
|  | Dark septate endophyte |  |
| | Total | $\frac{AMColonised + DSE + (AM + DSE)}{TotalRoot}$ |
| Ericoid mycorrhiza | Blue coils (Bl) | $\frac{BlueCoils}{TotalRoot}$ |
| | Brown coils (Br) | $\frac{BrownCoils}{TotalRoot}$ |
| | Type two | $\frac{TypeTwo}{TotalRoot}$ |
| | Dark septate endophyte | $\frac{DSE + (ErM.DSE)}{TotalRoot}$ |
| | Total ErM colonisation | $\frac{Bl + Br + TypeTwo + MixedErM + (ErM + DSE)}{TotalRoot}$ |

### Performance of the model efficacy with colonisation assessments

We quantified performance using an *F_1_* Score (see Eq. 1) which calculates the harmonic mean of *Precision* and *Recall* (see Glossary) yielding a value between 0 and 1, where 1 indicates the best performance (Chinchor 1992, Grandini *et al*. 2020, Reinke *et al*. 2021, Fergus *et al*. 2022). There are two equivalent definitions, either using *Precision* and *Recall* or the number of true positives (*TP*), false positives (*FP*) and false negatives (*FN*):

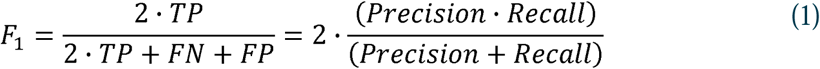

The *F_1_* Score penalises extreme values of either *Precision* or *Recall* and rewards models having similar high values for both, whilst simultaneously being robust against imbalanced datasets (Fergus *et al*. 2022, Khan *et al*. 2021, Crespo-Michel *et al*. 2023, Rahman *et al*. 2023). This was a key advantage over the *Accuracy* measure (see Glossary) originally used to quantify the predictive performance of AMFinder in Evangelisti *et al*. (2021). We applied an *F_1_* Score to each individual class and across the entire model as *Macro F*_1_ , thus incorporating corresponding *Precision* and *Recall* scores across all classes (Fig. S2) (see Eq. 2, Opitz 2024).

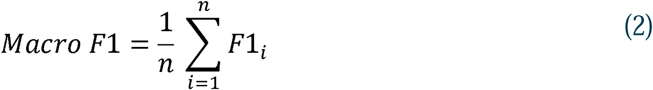

### Development of a user-friendly interface

To improve user experience and accessibility for non-computer experts we: 1) streamlined the way the user interacts with the application, by allowing the user to access all the tool functionalities including training via a single user interface (Fig. S3; see Methods S2 for further details on model training); 2) upgraded the user interface to encapsulate the end-to-end workflow in the tool, from uploading TIFF-format slide images and splitting them into smaller JPGs for utilisation by the tool to calculating predictions and outputting colonisation percentages (at the root level) and uncertainty measures, thus simplifying the workflow to allow all activities to be carried out via a centralised point of access and decision making; 3) added a confidence threshold setting to the existing workflow to ensure users can review uncertain predictions and overwrite them manually if needed. This modification ensured that there was a human in the loop for assessing the output of the machine learning model and enabled users to balance automation with certainty; 4) added high-level image overviews for reference while annotating and included the ability to save annotations and maintain unknowns temporarily (using ‘?’) without needing to close the tool. This function was intended for less experienced analysts allowing them to optionally embed comments and proceed with other annotations while awaiting expert supervisors input on uncertain tiles; 5) migrated the user interface from a standalone application written in the OCaml programming language to a more versatile web-browser-based application written in *React*; 6) replaced the data storage system of zipped CSVs used in AMFinder with a PostgreSQL database for storage of predictions and annotations, aligning with industry standards and improving data governance; and 7) added a persistent configuration to the tool to allow users to save settings between sessions (for example whether to use contextual mode when generating predictions).

## Results

### Quantification of AM fungal colonisation in field-collected roots using MF

Automated predictions of colonisation improved significantly compared with the original AMFinder tool as seen in the calculation of colonisation using the automated function (Fig. 2). By adding manual annotation to these (in tiles beneath the set confidence threshold), accuracy on par with microscopic examination was achievable.

**Figure 2.**
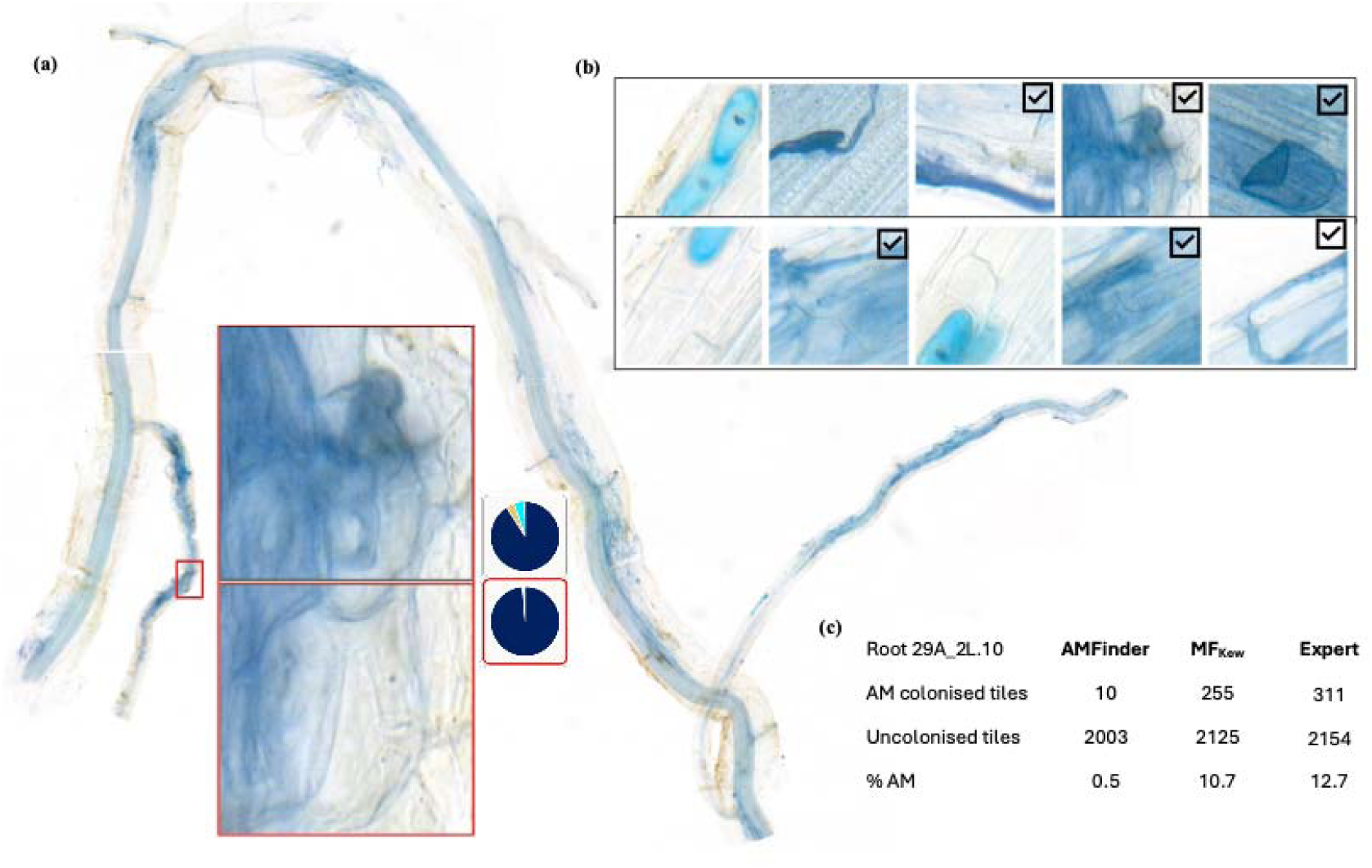
Comparison of colonisation predictions using the original AMFinder and MycorrhizaFinder (MF) on the same arbuscular mycorrhizal (AM) field-sourced root after clearing, staining and high-resolution scanning. a) Image root section (1 cm in length), showing example of two colonised tiles (red-lined boxes) that were not counted as colonised by AMFinder (i.e. false negatives) but were correctly predicted by MF. AM colonised is represented by dark blue in corresponding pie-charts. b) The ten tiles predicted as AM colonised by AMFinder, black tick boxes on 6 of 10 tiles indicate predictions are expert validated leaving 4 of 10 as false positives. c) Summary of tile counts and colonisation predictions generated by AMFinder versus MF for this root are significantly different. Human expert analysis of tiles is included for contrast.

AMFinder repeatedly misclassified field root tiles. In the example shown in Fig. 2b, four of the ten tiles predicted as AM^+^ were incorrect due to their biological ambiguities. Overall, MF achieved significantly better detection of AM^+^ class tiles resulting in prediction of 10.7% AM^+^ colonisation compared with 0.5% for AMFinder, before human augmentation and despite biological and technical artefacts e.g. patchy staining, soil debris. Expert estimates of colonisation were 12.7%.

### Incorporating contextual information significantly improves prediction quality

Tile-level predictions on an AM-colonised root region with and without the contextual function can yield significantly different determinations of class, affecting root-level colonisation percentages (Fig. 3). We observed a consistent improvement of the mycorrhizal class performance function when activating the contextual function, independent of the model architecture (Fig. 4). The *F_1_* Score for all classes improved with context using both the AM and ErM pathways (except for *DSE*), while the overall *Macro F_1_* improved by 5.0% for AM and 5.8% for ErM. Combining the model architecture and context function changes, the *Macro F_1_* improved significantly by 44.9% and 40.4% for AM and ErM, respectively, when compared against AMFinder (e.g. compare *Macro F_1_* in Fig. 4 with Fig. S1).

**Figure 3.**
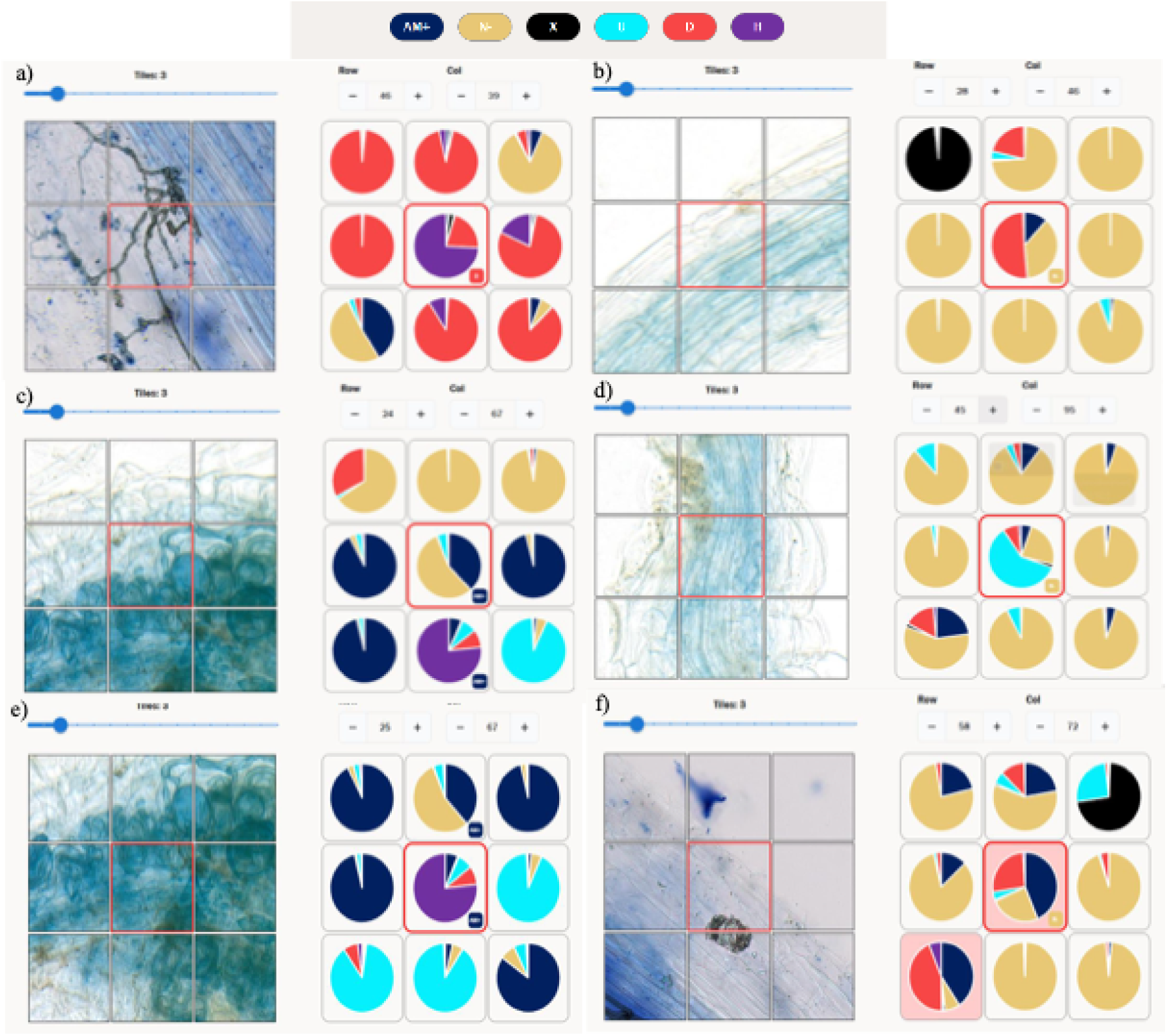
Examples of tile classes that switch when applying the contextual function in MycorrhizaFinder for the arbuscular mycorrhiza (AM) pathway. The class to which it changes is shown in the lower right corner of the central, red-bordered target tile. Tile class designation before and after context implementation for the six examples are detailed here: a) AM+DSE (H) to DSE (D); b) DSE to Uncolonised (N-); c) Uncolonised (N-) to AM Colonised (AM^+^); d) Unreadable (U) to Uncolonised (N-); e) AM+DSE (H) to AM Colonised (AM^+^); f) AM Colonised (AM^+^) to Uncolonised (N-). See Table 2 for tile class descriptions, and Fig. S3 for an illustration showing the extent of overlap by neighbouring tiles when activating the context function.

**Figure 4.**
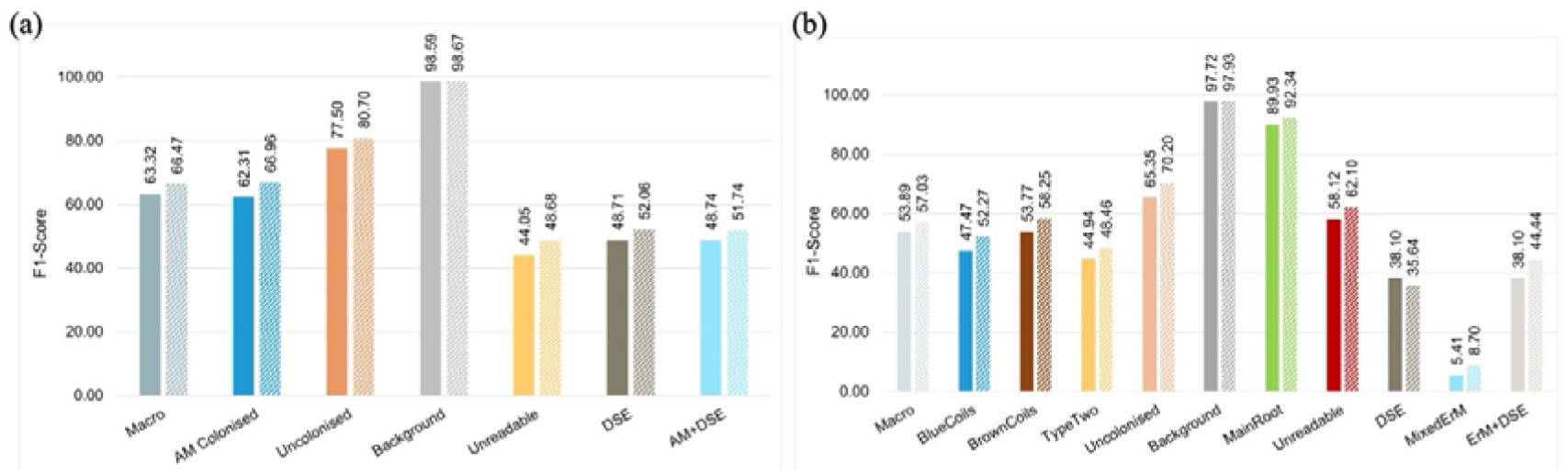
Performance (*F_1_* Score) of MycorrizaFinder_Kew_ with and without the context feature (lighter and darker paired colour bars, respectively). a) The arbuscular mycorrhiza (AM) path on our test dataset of AM host roots. b) The ericoid mycorrhiza (ErM) path on our test dataset of ErM host roots. *Macro* reflects the *F_1_* Score (Eq. 2) for the entire model averaged over all classes; the other bars refer to the individual *F_1_* Scores of the corresponding classes. Scores are expressed as a percentage, where the maximum score is 100. Scores have been obtained on the same model weights with all confidence thresholds set to 1, without any human supervision. See Table 2 for tile class descriptions.

In the AM case (Fig. 4a), the highest class *F_1_* Score was consistently *Background*, with values over 98 (both with and without context), followed by the *Uncolonised* achieving 77.5 (non-contextual) and 80.7 (contextual). These two classes were also the most represented in the training dataset (Tables S6-S9), with corresponding relative shares of 74.7% and 17.9%.

However, balancing measures applied in the pre-processing steps during the training stage reduced their actual number to the count of the *AM Colonised* “anchor class”, which the other model classes were measured against (Methods S2). With the ErM dataset (Fig. 4b), the highest *F_1_* Scores were observed for the classes *Background*, *Main Root* and *Uncolonised*, attaining 97.7, 89.9 and 65.4, respectively.

For both datasets and models, the application of the contextual function improved the *F_1_*Scores in most classes, with few exceptions. This also held true for the shallower CNN1 model architecture (Figs. S3a,b).

### The F1 Score provides a robust and interpretable estimate of colonisation accuracy

We found that the additional tile classes enabled better recognition of mycorrhiza structures, (see *F_1_* Scores, Fig. 4). Differentiating tiles with DSE and other artifacts typical of field roots allowed the machine learning to focus on the mycorrhizal classes.

Classes with higher sample representation yielded a higher contribution to the model *Accuracy* than classes with lower representation as seen in the performance of the ErM model (no context) for an independent test set of the classes ‘Unreadable’ and ‘DSE’ (Fig. 5). The *Macro F_1_* not only gave all classes the same weight towards a single performance measure (independent of their prevalence in the dataset) but also provides clarity on whether all samples for a given class were captured (*Recall*) and actually correct (*Precision*). While the *Accuracy_DSE_* was measured at 76.92 on the independent ErM test dataset, suggesting a high performance if this was the only used metric, the corresponding model *Precision_DSE_*was measured at 25.32, leading to a class *F_1_* Score*_DSE_* of 38.10. Inversely the *Accuracy_Unreadable_* of 52.50 suggested lower performance, if it was the sole component of the metric. The corresponding *Precision_Unreadable_*of 65.09 raises the total *F_1_*Score*_DSE_* to 58.12.

**Figure 5.**
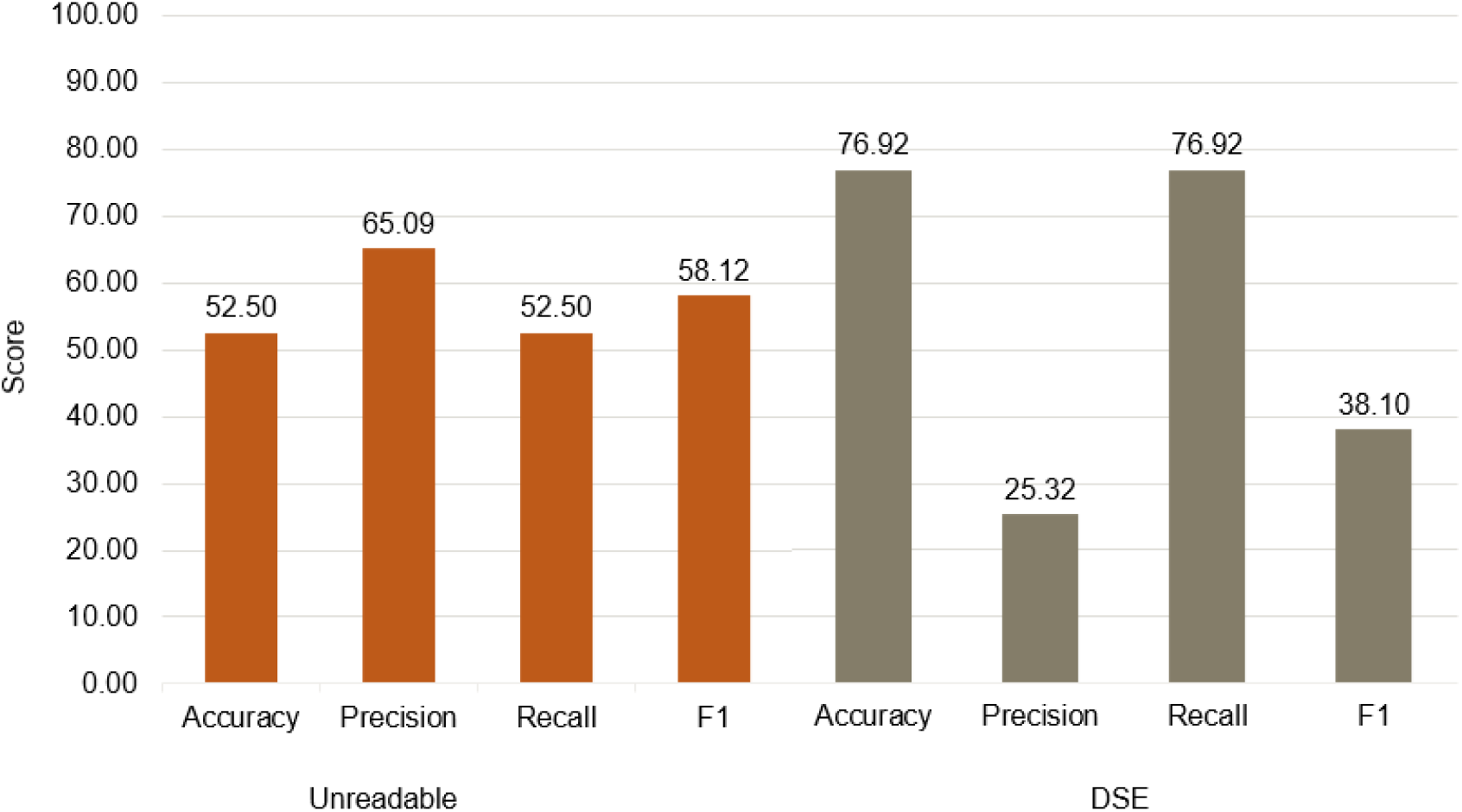
Quality metrics for the underrepresented classes ‘Unreadable’ and ‘DSE’, as measured on an independent test set for the ErM data and using the deployed EfficientNetB5 model (no context). Calculations of *Accuracy* and *Recall* for the Unreadable and DSE classes are the same as the denominator, in this case is the same. See Table 2 for tile class descriptions.

### MF reliably facilitated accurate quantification of ErM fungal colonisation in field-collected roots

The ErM pathway was able to automatically distinguish features across the ErM classes *Blue coils*, *Brown coils*, *Type two* and *MixedErM/ErM+DSE* most of the time, and performed better with context. Representative examples of the tool distinguishing amongst biological features and artifacts (without human corrections) are shown in Figure 6. The central tile (Fig. 6a) predicted Bl+ while the tile above initially predicted HE+, but with context, corrects to Bl+. In Figure 6b the model predicted the central tile Br+, with high confidence, while the tile to its right is predicted U. The lower row yielded two of three tiles with pink backgrounds showing that the threshold of certainty for all potential classes was below the conversion threshold (here set to 0.5). This shows how the programme signals to the user that a manual annotation is required if the tile is to be included in the calculation of colonisation metrics. In Figure 6c the panel shows the ability of the model to distinguish the thicker T+ hyphae from the more common Bl+ coils.

**Figure 6.**
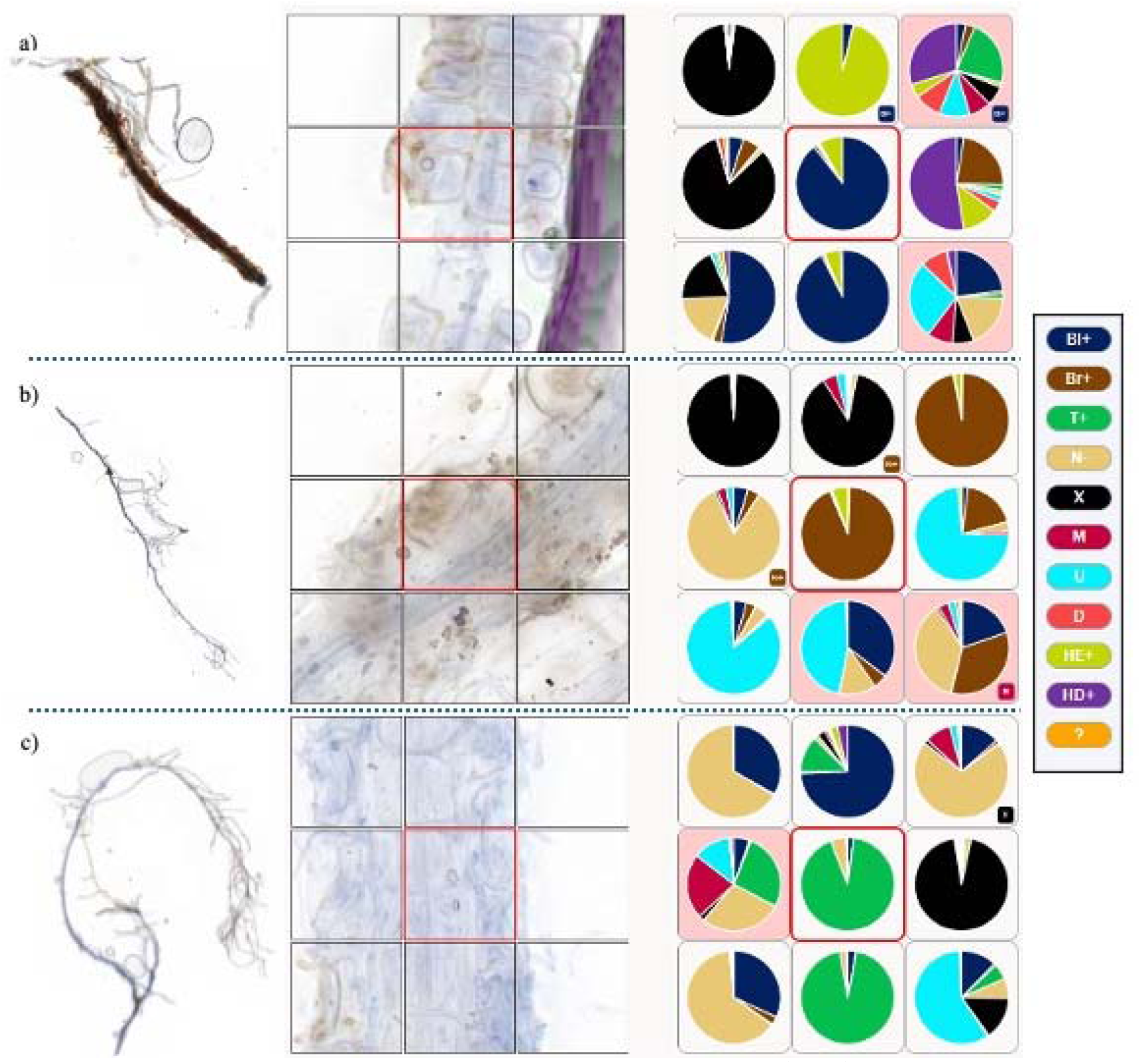
Representative images of three ericoid mycorrhizal (ErM) field roots and class predictions using the MycorrhizaFinder ErM pathway. Central images show a nine-tile panel magnifying a section of root. The pie-charts show corresponding class predictions, with the contextual function switched on. The legend details one of ten class possibilities (see Table 1 for reference of tile class abbreviations). The central, red-bordered image tiles were predicted as a) Blue coils (Bl+); b) Brown coils (Br+); and c) Type-2 patterns (T+). A small box label in the lower right corner of the classification tile indicates that the context function yielded a different outcome and amended the tile prediction. A classification tile with a red background indicates that the predictions were below a set conversion threshold (default 0.5) pointing the user to manually annotate.

A dedicated ErM model path significantly improved the biological ambiguity in distinguishing ErM septate hyphae structures from AM aseptate hyphae in roots from complex field environments. With the AM path, ErM structures were not typically misclassified as AM fungi, but commonly misidentified as DSE, thereby underestimating mycorrhizal colonisation (see Table S5). Our model development showed that with more training data, the individual *F_1_* Scores continue to improve.

### Throughput was faster and simpler

Automated predictions of root colonisation using MF, as with AMFinder, significantly reduced analytical time compared to traditional techniques involving a human using a microscope. But with MF, augmentation of classes by user supervision was greatly simplified by allowing modifications of tile classifications within the same interface, without exiting and reloading, which was necessary previously, and single root images are now loaded in one step without splitting into several JPGs (see Fig. S6) as the tool can now manage significantly higher resolution jpgs than previously. Moreover, if necessary, conversion from TIFF to JPGs was also supported by the tool interface. Finally, the user no longer required programming skills to start-up the tool and could run it on a laptop on Windows, MacOS or Linux, albeit this is slower than using a dedicated desktop computer.

### Performance using other image capture devices

The MF model performed well on a subset of images (from the Evangelisti et al. 2021 dataset) captured using a high-resolution light microscope. The 3-class *Macro F_1_* was 58% (without enhancements) compared with our trained roots of 65% (Fig. S7). Previously, the model was wrongly predicting tiles containing AM vesicles and hyphae, or uncolonised cells as DSEs (Figures S3, S4).

### CNN1 model vs EfficientNetB5

The EfficientNetB5 architecture (Tan and Le, 2019) achieved high classification performance on a fully independent test dataset while being operational and retrainable on local GPUs (Table 4) and was ultimately chosen for the final tool deployment. Larger models, such as EfficientNetV2M (Tan and Le, 2021) did not achieve significantly higher performance despite being built on a significantly higher number of parameters (EfficientNetB5 at ∼30M vs. EfficientNetV2L at ∼54M, Table S1).

**Table 4.**
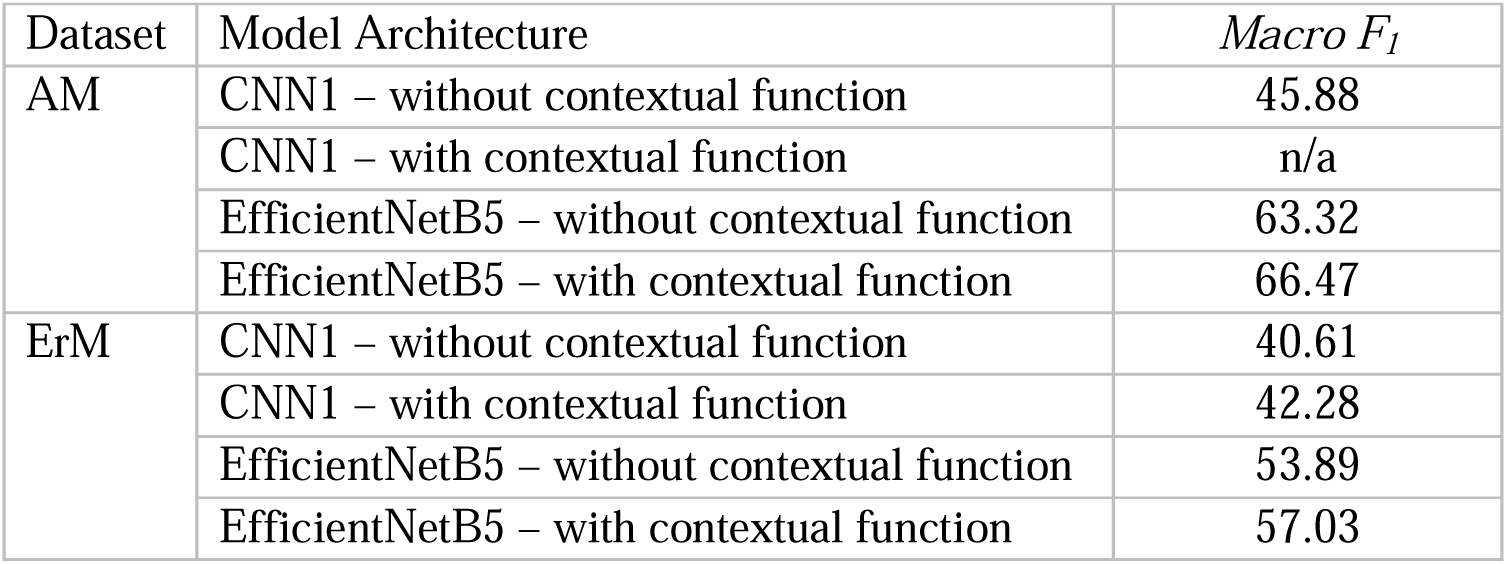
*Macro F_1_* Scores for individual EfficientNetB5 (Tan and Le 2019) model architectures alongside the best CNN1 models as a benchmark comparison for the Arbuscular (AM) and Ericoid (ErM) Mycorrhiza datasets. The contextual function was not implemented for the AM CNN1 model.

There was a significant improvement of the *Macro F_1_* Score when comparing the simpler CNN1 model to the EfficientNetB5 model for both the AM and ErM pathways (Table 4). The EfficientNetB5 model achieved higher classification performance when trained on the same dataset. Since both model architectures were optimised on the same training dataset, we concluded that the deeper model architecture of EfficientNetB5 captures complex image features in the dataset more effectively, leading to higher classification performance. This was also observed when comparing the individual class *F_1_* Scores achieved in the CNN1-model with the ones obtained on the EfficientNetB5 model (compare Figures S7, S8 with Figure 4).

## Discussion

Standardised, medium to high throughput methods for mycorrhizal root colonisation assessments in field roots are needed by researchers, policy-makers and land managers to provide repeatable processes to monitor environmental change across scales. Such a method will facilitate direct comparison across different studies and ecosystems. Our new MF tool addresses a research gap by presenting a computer vision tool specifically trained and optimised for field-collected roots successfully differentiating common artifacts and disturbances arising from field collection with different endomycorrhizal types (through the AM and ErM pathways) It has also significantly improved access for researchers to assess root colonisation for controlled glasshouse and laboratory experiments. in a user-friendly interface.

By applying multiple technical innovations to AMFinder, our upgrades provide an enhanced user experience with an intuitive design. The portable *React* (Meta Open Source 2025) implementation results in a tool accessible to non-experts and adaptable across all major computer operating systems. The user interface allows human oversight on the predictions calculated by the model. Thus, expert tasks are reduced to assessing the machine predictions of the relevant tiles of each microscope image, with confidence thresholds determined by the user, thereby inviting trade-offs between certainty and time inputs root by root. The tool can be run on a single desktop computer without requiring advanced hardware, though prediction time is reduced when a dedicated GPU is available. Replacing the preexisting CNN1 model with a pretrained EfficientNet (Tan and Le 2019) architecture, allowed us to introduce new classes for AM assessments - *DSE,* mixed *AM+*DSE and *Unreadable* - to improve the performance of assessing colonisation by removing ambiguity in visual recognition. The higher classification performances of *Background* and *Uncolonised* was mainly due to less intricate image features compared to the other classes. Experiments and performance measures obtained on corresponding CNN1 models outlined a comparable classification behaviour (compare with Figs. S7, S8). The model’s lower *F_1_* Scores in the cases involving classification of fungal structures (*AM Colonised*, *DSE Colonised* and mixed *AM+DSE Colonised*) signify higher complexity in image features compared to the *Background* and *Uncolonised* classes. Taking the relatively constrained shares of data into account (Table 1) i.e. 0.9% for the DSE *Colonised* class and 0.3% for the *AM+DSE Colonised*, the obtained *F_1_* Score result was analytically meaningful and indicates the model architecture’s capacity to generalise. A similar reasoning could be applied when revisiting the results of the *Unreadable* class, adding that the image features encountered in this class were based on blurry, out-of-focus or indefinable diversity instead.

Our new model for ErM colonisation arose because the AM pathway, trained on aseptate fungi, is not suitable for recognising septate ericoid hyphae classes as evidenced by the example shown in Table S5. By training a separate path for Ericaceae roots, we are able to more efficiently evaluate Northern European Ericaceae-dominant habitats at scale. In most cases, the sample collector will know if the host is Ericaceae as these woody shrubs are apparent year-round. The model’s *Main Root F_1_* performance is stronger than the ErM classes, providing confidence that the model is already able to distinguish the older more lignified root tiles leading to a more reliable estimate of colonisation. These values could be attributed to a similar rationale as in the case of the AM dataset i.e. image features encountered in the *Background* and the *Main Root* classes were more limited in their diversity and were easier to classify. A higher relative share in the training data could be excluded as these classes originally exhibited a higher level of representation compared to the *Blue Coils* “anchor class” and were thus balanced via clipping (Table S3, Methods S2). This was further supported by results obtained on the CNN1-ErM model (see Fig. S3b), where a comparable distribution in classification performance could be observed. The classes that characterise the colonisation of Ericaceae roots - *Blue Coils*, *Brown Coils*, *Type Two*, *DSE*, *Mixed ErM* and *ErM+DSE*, could be clustered into two performance groups. The first three achieved more robust *F_1_* Scores around 50.0, which could be attributed to higher relative data shares (2.8%, 3.0%, 0.4%, respectively), and the latter three demonstrated greater variation in the obtained *F_1_* Scores, which could also be attributed to relatively low representation in the training dataset (2.3%, 0.4% and <0.1%, respectively). It appears however that the *ErM+DSE* class exhibited more recognisable image features compared to the *MixedErM* class, as a higher *F_1_* Score was achieved despite significantly fewer images present in the data.

Given phenotypic complexities of ErM in field roots, by introducing new tile classes distinguishing different morphotypes within ErM (distinctly blue or brown colours and Type two), we aimed to be able to recognise potentially ecologically relevant signals occurring at the habitat scale. We were unable to conclusively attribute the blue and brown putative ErM to hyphal ontogeny or staining artifacts, especially given extensive staining trials of *Calluna vulgaris* roots using pen ink and Trypan blue stains at the individual root level. Regardless, the ErM types can easily be aggregated together by the researcher, and we do not find any drawbacks in the model performance. Recognising DSE hyphae on their own, but also when co-occurring with either ErM or AM, the trained model can better differentiate the anchor mycorrhizal classes. It also facilitates better understanding of DSE prevalence across field samples and specific habitat types, while simultaneously ensuring that the mycorrhizal classes are not underrepresented when they co-occur within a tile. There are still biological ambiguities to unpack, e.g. compressed vascular bundles mistaken for DSE hyphae, morphological variances arising from rare plants or habitats, but if the user also learns to spot these types of errors, they can easily resolve them. The contextual information added to the model by incorporating the predictions for the surrounding tiles into each tile’s classification has also boosted model predictions significantly and improved F_1_ in most classes. Negative trends were notable for the classes Type 2 and DSE in the ErM dataset. This was likely due to the low sample sizes for these classes, but we maintained them in the training set to support more clarity on ErM colonisation classes. As the contextual classification exploits max-voting, this could lead to other classes outnumbering the image tile with said colonisation and thus a false relabelling.

The resulting MF tool performed well overall and was deemed successful in its capacity to improve the pace and volume of colonisation assessments for field roots in our lab. Previously, a single root section would take half to one hour to evaluate using a microscope. Not only did this limit the number of roots per day per analyst to an average of ten, but the reproducibility under traditional microscopy methods has considerable challenges, e.g. complex root architectures make stopping and restarting in position difficult. The model can predict significantly more roots overnight (dozens to hundreds based on the hardware used). Depending on the confidence threshold designated by the user and the complexity of the root image, the expert analyser can refine these predictions to annotations in less time than traditional microscopy assessments. The tool outputs and database provide repeatability and accountability in tracking and storing the assessment scores and create a new reference visual database of roots for researchers and policy makers alike. This is a significant step towards enabling landscape-scale assessment of *in situ* mycorrhizal colonisation. However, we suggest that the resulting colonisation percentages are used as a starting point and considered within the context of experimental limitations (e.g. sampling size) and research questions.

Despite the output precision, categorical assessments may be more appropriate, e.g. a DAFOR scale (Sutherland 2006) which roughly delimits Dominant (e.g. >50% of root length), Abundant, Occasional, Frequent or Rare, particularly in real world settings.

Interpreting a percent colonisation in a controlled laboratory experiment with a known mycobiont is significantly different than in roots sampled to represent change at the landscape or habitat scale. Given the link between root mycorrhizal colonisation and functional outcomes for host plants, and the responsiveness of root associated fungal communities to environmental change (Soudzilovskaia et al. 2015), the tool has potential for supporting the use of mycorrhizal colonisation data as a contributing indicator of ecosystem condition and for natural capital assessments (Frew 2025). When coupled with an efficient low-cost staining protocol, it provides a viable option for medium-throughput colonisation assessment of field-collected roots for the first time.

The requirement for expert human intervention is a double-edged sword. While it can reduce the time saving advantage of the tool, and the potential users to those capable of identifying mycorrhizal structures within roots, it allows deep engagement and discovery in unravelling complexities at the cytological level, particularly useful when coupled with molecular data. We recommend a workflow which starts by setting a threshold value of 80% - 90% to allow expert time to manually focus only on tiles yielding unacceptable certainty. Future model development should focus on improving the predictive performance of the underlying machine learning model by targeting the classes that are influencing variability around estimations of AM or ErM colonisation. Since we observed a trend of improved predictive performance when adding more training data, a priority can include the labelling of additional training data especially in the underrepresented classes, e.g. ErM and DSE. Exploration of alternative model architectures such as vision transformers may enhance performance (noting that this will probably involve a trade-off with speed and computational requirements).

As there is no clear industry standard for performance-related metrics in image recognition tools, we selected a *Macro F_1_* Score as the main model performance metric, because it performs well on imbalanced datasets, acknowledging multiple approaches (Opitz, 2024), and represents a more holistic measure than other common metrics such as *Precision*, *Recall* or *Accuracy*. If we were to have reported test *Accuracy*, strictly defined, it would have been 91.55% for the 6-class EfficientNetB5 AM model (82.13% on the 3-class CNN, before upgrading) and 86.68% for the 10-class ErM model, but this risks overstating performance overall across classes. We therefore propose future model developments use these baseline *Macro F_1_* Scores to measure performance improvement. Although *Accuracy* is a metric that can be understood intuitively, it does not provide insight into the model performance across individual classes, especially needed with the evolved MF model given its imbalanced tile class structure (Grandini *et al*. 2020, Opitz 2024).

Importantly, because MF is published under an open government licence, it offers strong potential for community-driven improvement. While the published version of the tool is trained on a dataset diverse enough to provide a baseline to train and evaluate most Northern

European systems, it further offers the capacity to allow researchers with a more specific study focus (by habitat, host plant, etc.) to add their own training datasets and create their own versions of the tool which are likely to be more effective for their specific system. The intuitive user interface means that this can be done without any coding or AI knowledge by uploading jpg root images and following the User Documentation. For users with programming skills, the open-source code on GitHub enables extension of both front-end and back-end functionality. This openness encourages collaborative development, accelerating model performance improvement and broadening MF’s applications, ultimately supporting high-throughput assessment of mycorrhizal colonisation across diverse ecological and practical contexts.

## Conclusion

Unsupervised, MF reaches acceptable mycorrhizal colonisation assessments across plants, fungi and habitats, especially differentiating *Uncolonised* tiles from other colonisation classes. By adding active expert human supervision, MF enables remarkable accuracy, especially with thoroughly-stained intact root. It provides users having limited prior knowledge of training machine learning models with a simple workflow to train with new image datasets and for the user to add tags to tiles further differentiating morphological types within classes. Beyond the innovations for controlled experiment root assessments, the tool’s adaptability to field-sourced variation in AM-host root biology, mycorrhizal structures and miscellaneous artifacts, and the introduction of a new model to assess habitats dominated by ericoid mycorrhizal fungi and differentiate dark septate endophytes, are of important value to researchers interested in monitoring and studying mycorrhizas in nature.

## Acknowledgements

Some of the arbuscular mycorrhizal roots used to train the model were collected from the Coronation Meadow at Wakehurst, under Kew’s Nature Unlocked programme. Staff within Kew’s Mycorrhizal Ecology laboratory provided time and resources to ensure the success of this project. We appreciate the labelling of Ericaceae roots by Ruairi Hafferty-Hay under the Air Pollution Recovery Programme funded by the Joint Nature Conservation Committee and we thank Ellie Wilding for preparing Figure 1. Finally, we acknowledge the thoughtful support of Rupert Thomas (Cambridge Consulting) with initial efforts on the tool’s development.

## Author Contributions

JK, LMS, AK, MB conceived of the modified and enhanced MF machine learning tool; RU, SB, JK, JS, KPT, ES, MR tested and trained the model and related user codes, streamlined the tool user interface and updated the model architecture and curated the root images. SS & EE provided substantial insights based on their experience developing AMFinder and validated the tool. All authors provided valuable and critical feedback and participated in the writing of this manuscript.

## Data Availability

MF is publicly available under an MIT licence. Executables for Windows, MacOS and Linux are available alongside comprehensive documentation at https://doi.org/10.5281/zenodo.17979413, the images used as training and test data are available at https://doi.org/10.5281/zenodo.17476686 and the source code is available at https://github.com/MycorrhizaEcologyLab/mycorrhiza-finder/tree/v5.0.0.

