## Supplemental Information for "MycorrhizaFinder: a user-friendly machine learning tool to quantify endomycorrhizal colonisation of real-world roots"

^1^ Ecosystem Stewardship, Royal Botanic Gardens, Kew, England; ^2^ Natural History Museum, London, England, ^3^ Imperial College London, England; ^4^ Institut Sophia Agrobiotech, UMR INRAE 1355, Université Côte d’Azur, Sophia Antipolis, Provence-Alpes-Côte d'Azur, France; ^5^ Sainsbury Laboratory, University of Cambridge, England; ^6^ d-fine GmbH, Germany; ^7^ Institut Botànic de Barcelona (IBB), CSIC-CMCNB, Spain.

Corresponding Authors: Jill Kowal and Laura M. Suz

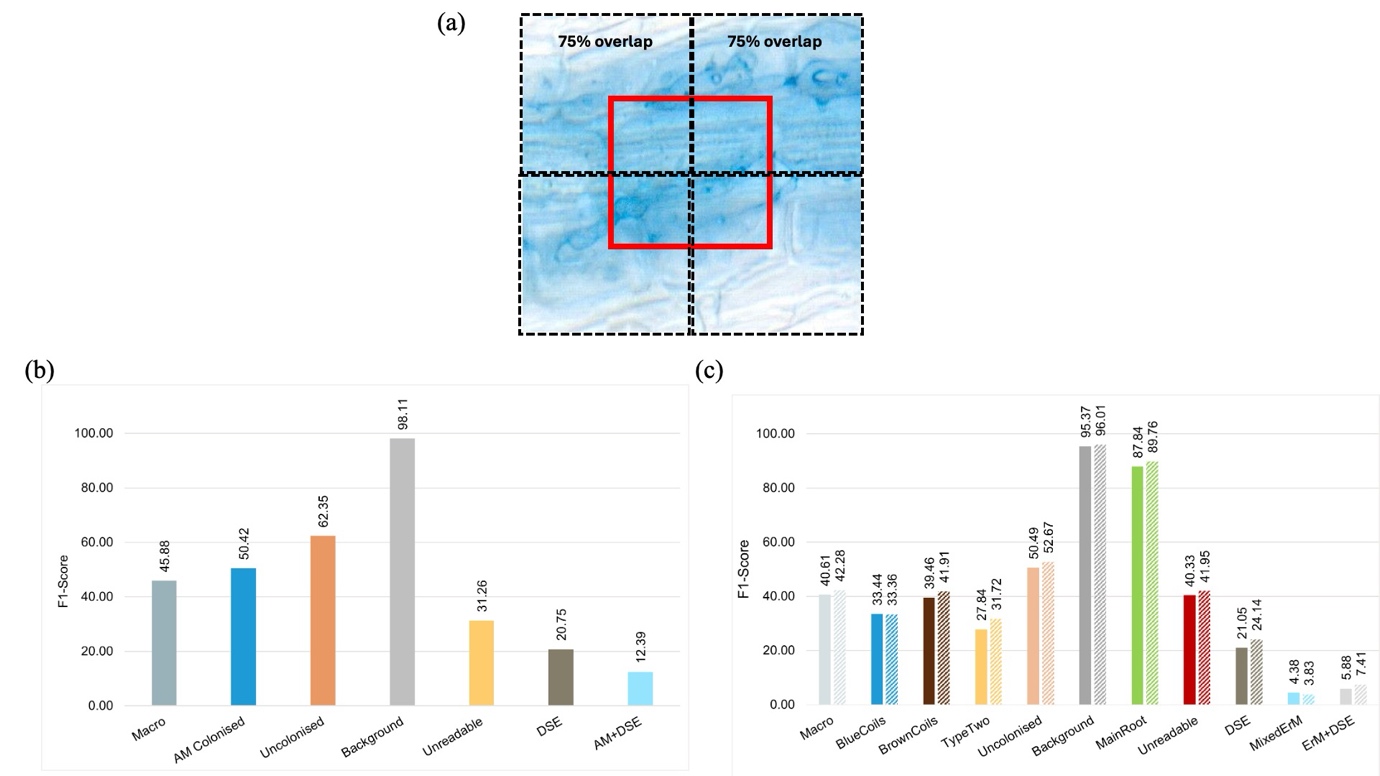
**Figure S1.** Contextual function effect on performance (*F_1_-*Scores). (a) Image of a tile (red outline) showing the extent of the context function covering four neighbouring tiles (outlined with dashed lines), overlapping by 75%. Performance using the original CNN1 model on the (b) arbuscular mycorrhiza (without context only); and (c) the ericoid mycorrhiza (without and with context, darker and lighter bars, respectively) datasets; see Figure 4 to compare CNN1 results with EfficientNet. *Macro* reflects the *F1* Score (Eq. 2) for the entire model averaged over all classes, while the other bars refer to the *F_1_*Scores (Eq. 1) of the corresponding classes. Scores are expressed as a percentage, where the maximum score is 100. Darker (left) bars within each class refer to the performance when ignoring context (of four overlapping tiles) by treating each tile independently, while right bars show the performance when accounting for context. Scores have been obtained on the same model weights.

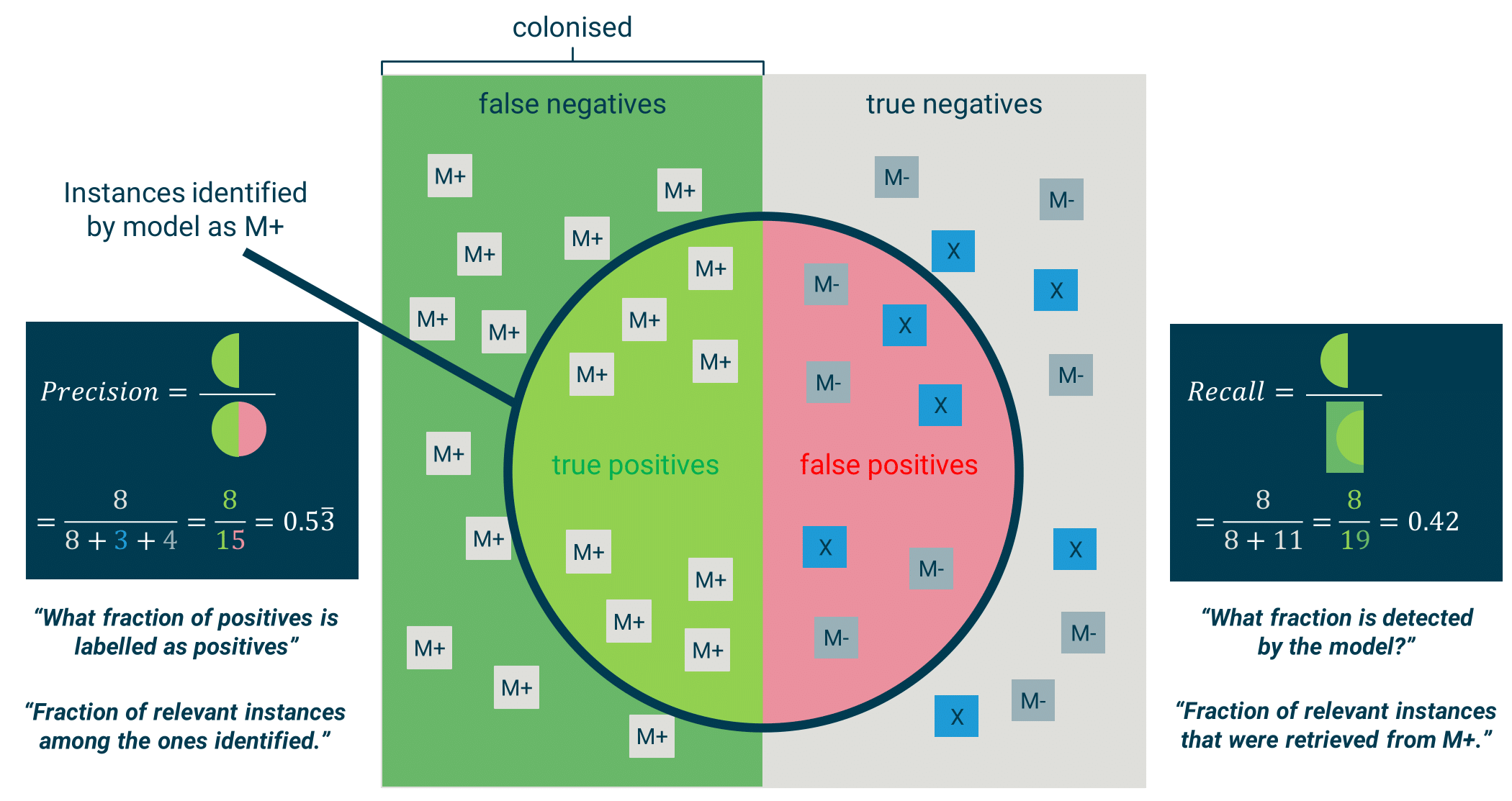

**Figure S2.** Graphical illustration of the quantities *Precision* and *Recall*, using a simplified scenario where the model is distinguishing between three classes: colonised (M+), uncolonised (M-) and background (X).

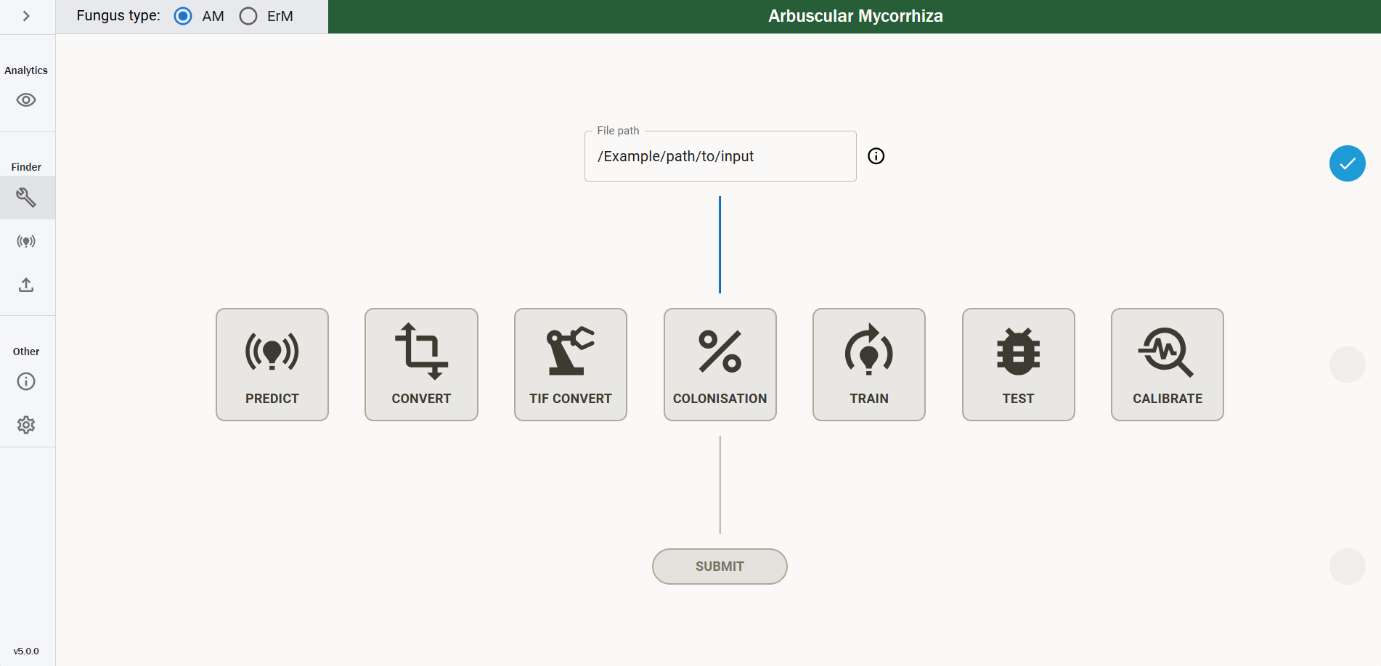
**Figure S3.** Main user interface panel of MycorrhizaFinder showing the functions accessible in the integrated platform: *Predict*, *Convert* (to colonisation metrics), *TIF Convert* (to JPG image format), *Colonisation*, *Train*, *Test*, *Calibrate*.

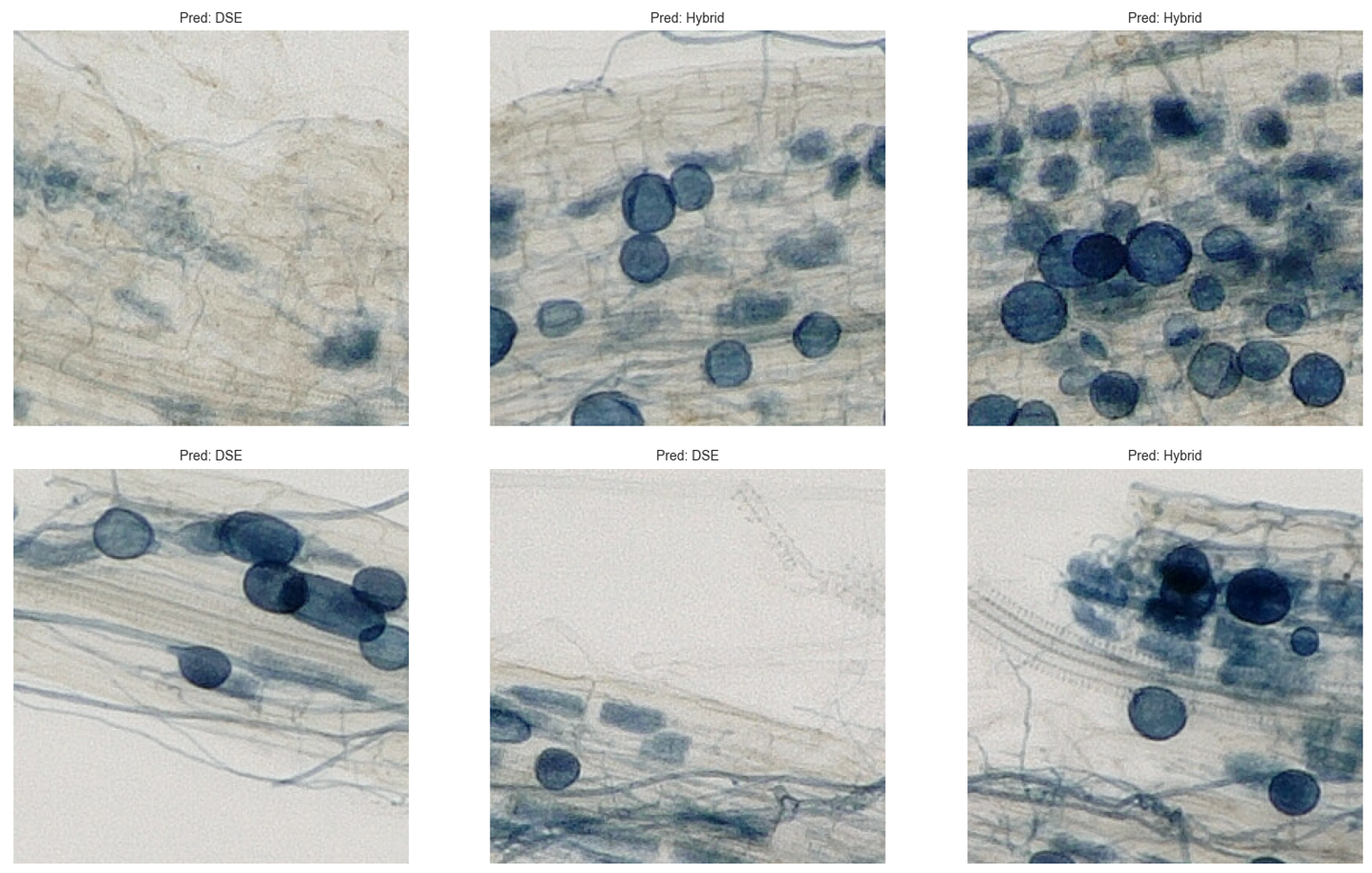

**Figure S4.** A subsample of AMFinder image tiles (from Evangelisti et al. 2021) labelled as *Colonised* (the equivalent of *AM Colonised* in MycorrhizaFinder (MF) due to the arbuscular mycorrhiza (AM) intracellular vesicles, arbuscules or hyphae, but incorrectly classified by the MF model. The MF model is sometimes incorrectly identifying AM hyphae as dark septate endophytes (DSE), leading to a classification of *DSE* or *AM-DSE*.

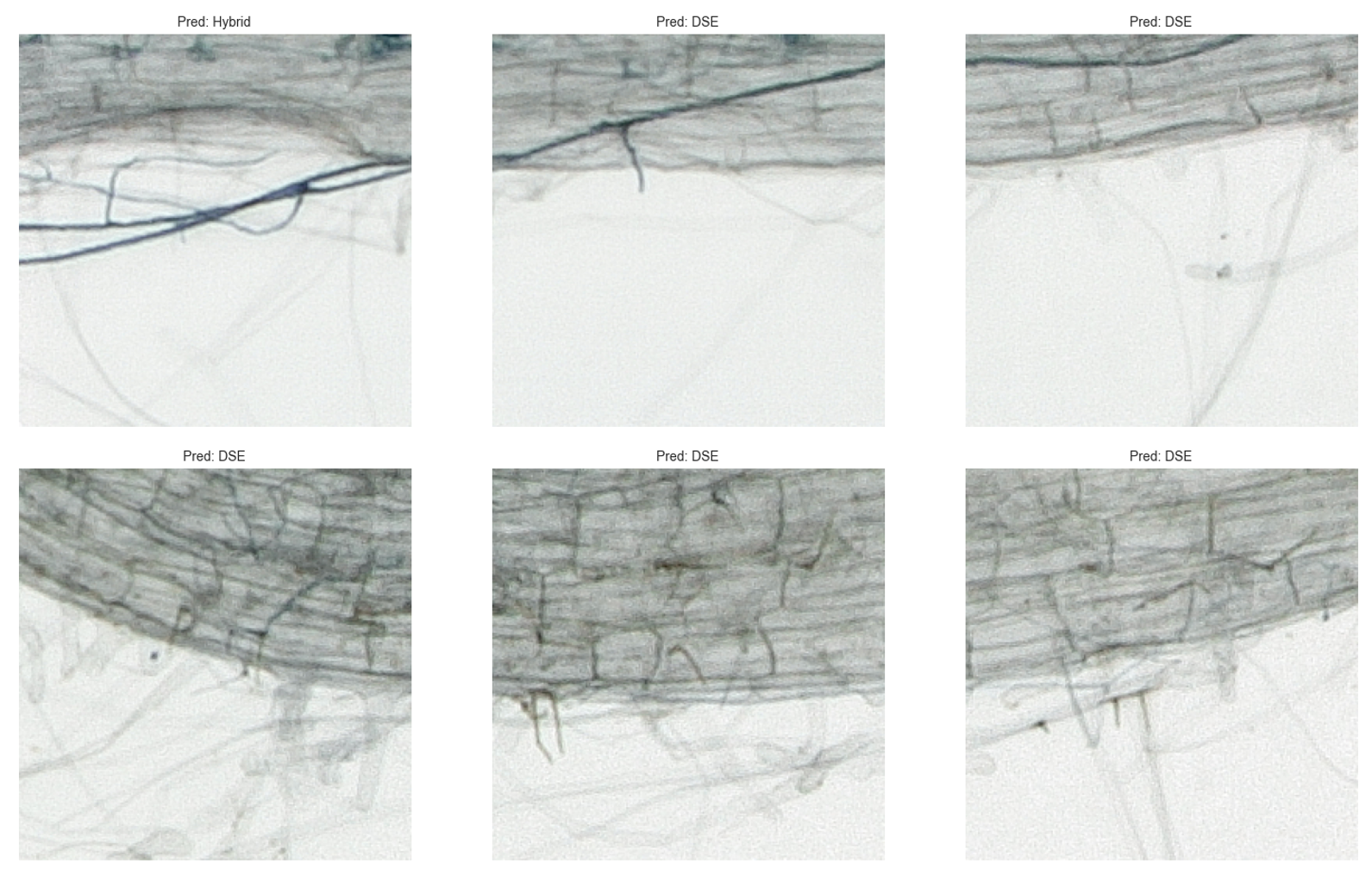

**Figure S5.** A sample of images from Evangelisti et al. (2021) labelled as *Uncolonised*, which were incorrectly classified by the MycorrhizaFinder model. As in Fig. S4, the model is incorrectly identifying dark septate endophytes (DSE) in the image.

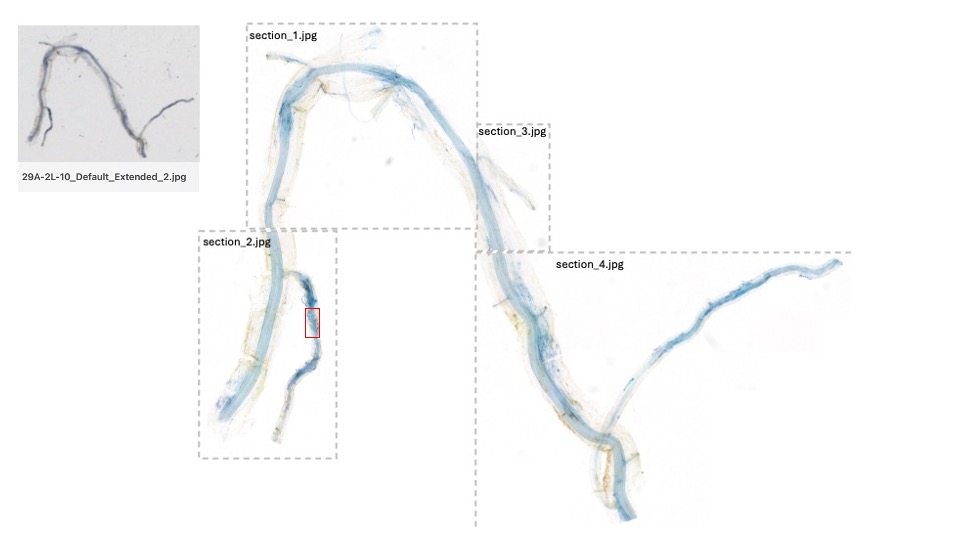
**Figure S6.** Image showing the extra technical step requiring partitioning of images into several jpgs (up to 16, depending on pixel sizes) before uploading to AMFinder. In order to generate colonisation predictions, each field root segment is analysed separately in AMFinder. Upper left image is 1cm long root on slide.

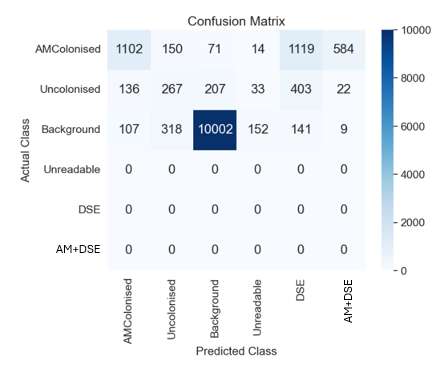

**Figure S7.** MycorrhizaFinder performance (shown in a confusion matrix) on a test using a subset of AMFinder (Evangelisti et al. 2021) trained images (tile sizes 252 x 252 pixels). The 3-class *Macro F_1_* was 58% compared with our trained roots of 66%.

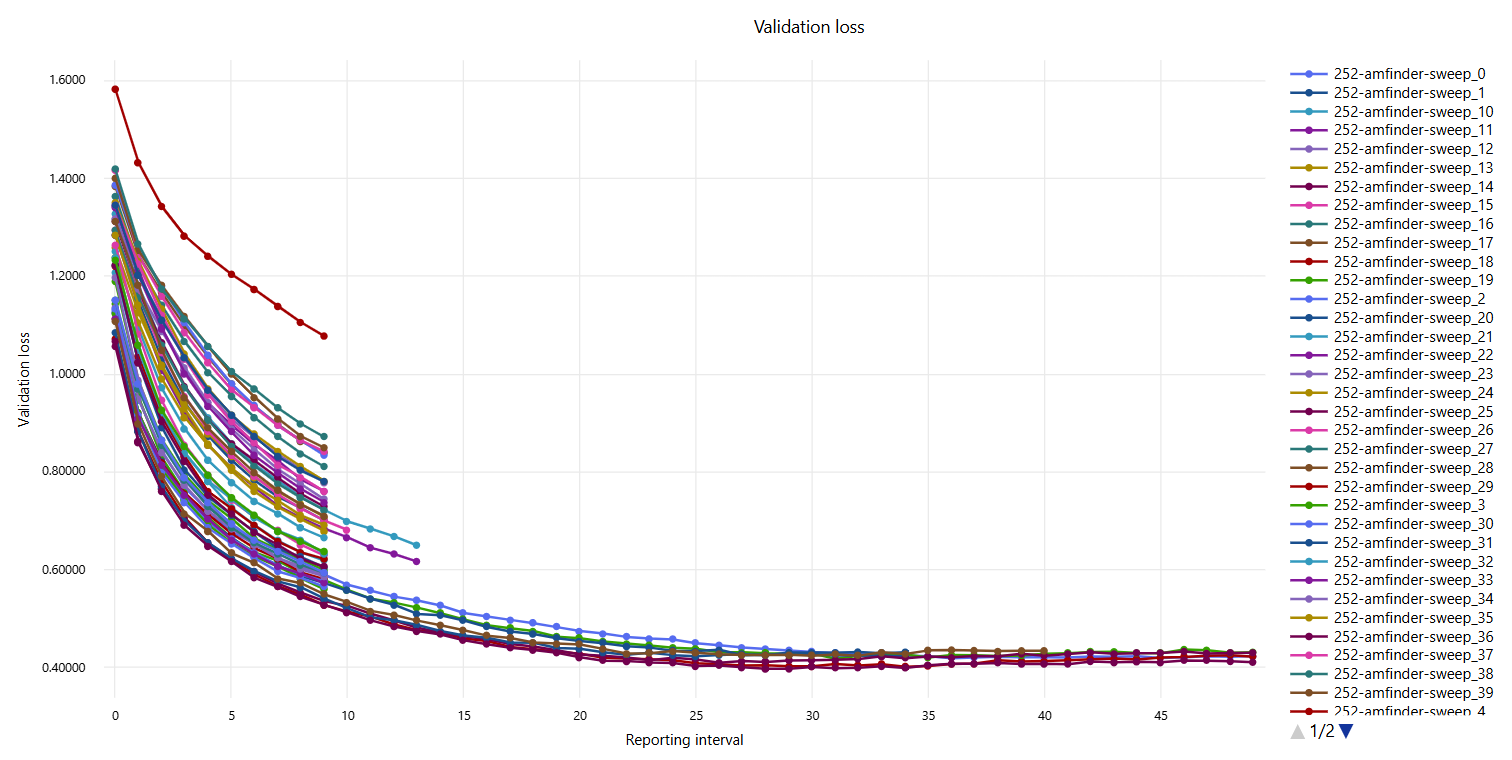

**Figure S8.** Exemplary hyperparameter sweep trials to train an arbuscular mycorrhiza EfficientNetB5 model. Trials that did not achieve a lower validation loss than the (then) leading trial, were truncated from the sweep after 10 epochs and reevaluated every five epochs.

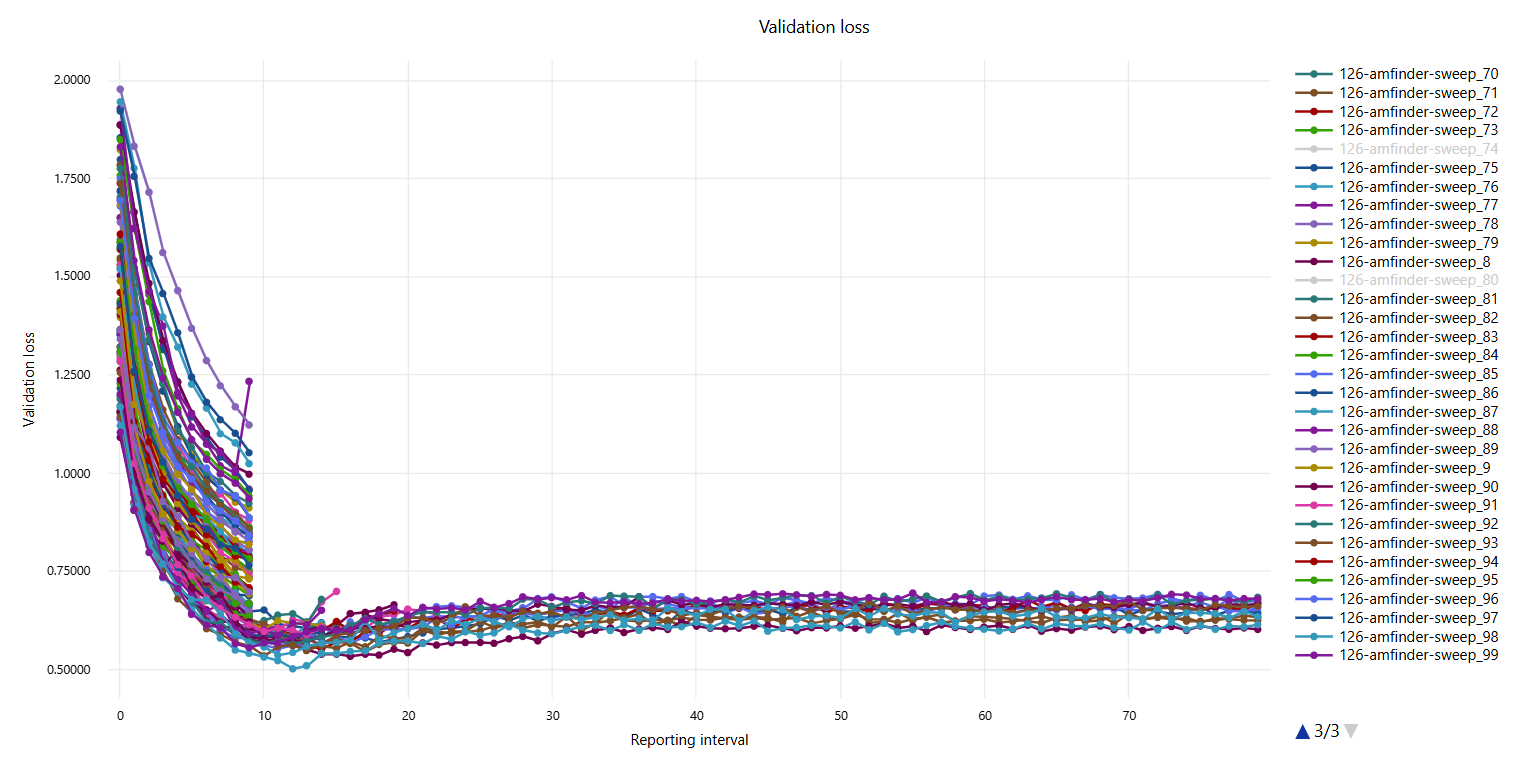

**Figure S9:** Exemplary hyperparameter sweep trials to train an ericoid mycorrhiza EfficientNetv2M model. Trials that did not achieve a lower validation loss than the (then) leading trial, were truncated from the sweep after 10 epochs and reevaluated every five epochs. Note that three trials have been excluded from the view to improve readability of other trials.

**Methods S1**. Kew’s modified staining protocol, adapted from Kowal *et al.* (2020) [10.21769/BioProtoc.3786](https://doi.org/10.21769/BioProtoc.3786)

Pre-staining

I. Set up two-well water bath to preheat, one to 60°C and the other to 100°C. (If one bath, start with the 100°C)
II. Take cleaned roots stored in 70% ethanol and select roots to stain. Try to separate roots that are visibly lignified (woody). For the staining of lignified roots, see modifications at the end of this protocol.
III. Transfer the roots to a 12-well or suitable staining dish for rinsing before staining – one root per well. Ensure that the wells are numbered and correspond correctly to the root code or sample name. At this point, the roots can be photographed if needed and cleaned carefully with a paintbrush if there is any visible dirt on them.

Root clearing

I. In the 12-well dish, rinse each root three times using dH₂O, ensuring each root is fully submerged in dH₂O during each rinse. Use a 1000µl pipette to remove the dH₂O, taking care not to draw up the root into the pipette tip.
II. Label 2ml new tubes (depending on the number of roots being stained) and add 500µl of 10% KOH to each tube.
III. Once rinsed, transfer each root to the appropriately numbered KOH tube and transfer the tubes to the 100°C water bath in a floating rack, ensuring that the lids do not rest below the surface of the water. Use cap locks to prevent the lids from opening while in the bath.
IV. Leave the tubes in the water bath for 20 minutes. After removing, check the tubes for any colour leached into the KOH. In cases where the KOH is particularly dark (dark yellow, orange, or brown), replace the KOH in the tube with fresh reagent. Repeat that step whenever you notice the KOH darkening.
V. Move the KOH tubes to the 60°C water bath for a further 25 minutes. After, check the roots and any that have cleared sufficiently (lignification removed and significantly paled in colour) can be carried forward to the next stage; any that still have lignification/ significant colouration can remain in the 60°C either until cleared or until the end of the day, checking on them periodically to assess both the root and the colouration of the KOH. If there are still dark roots after this, they can be left at room temperature over night to clear further.
VI. When roots are clear, remove the tubes from the water bath and return to the numbered wells of the 12-well dish.

Staining

I. In the 12-well dish, rinse each root three times using dH₂O, ensuring each root is fully submerged in dH₂O during each rinse. Use a 1000µl pipette to remove the dH₂O, taking care not to draw up the root into the pipette tip.
II. Any roots that are slightly yellow, or have some areas with pigmentation, can be submerged in 2% HCl prior to staining to bleach them. Leave these roots in the HCl for 2-5 minutes (you should see a colour change almost immediately).
III. Label fresh 2ml tubes and fill them with 500µl staining solution (see recipes for reagents).
IV. Gently transfer each root to the appropriately numbered staining tube, making sure they are completely submerged. Roots will be fragile after clearing, so transfer with care. Where possible, use the dissection needle rather than tweezers.
V. Place the tubes in the water bath at 100°C for 1-3 minutes depending on how thick the root is or how lignified the root was. Alternatively, if roots are very fragile, leave in ink-filled tubes on bench at room temperature for about 2-3 hours or trial with lower bath temperature for 3-5 minutes. Note that thicker or more lignified roots usually need less time as they absorb stain more readily and clearing them can be problematic.
VI. Remove the tubes from the water bath.

De-staining

I. Fill each well of the 12-well dish with 1% acetic acid.
II. Use the staining needle to remove roots from their respective tubes by gently scraping the tip of the needle along the bottom of the tube and up the side. The root should be visible as a dark blue string. Place the roots in their respective wells filled with 1% acetic acid to leach excess stain from the root.
III. Using a 1000µl pipette, gently remove the acetic acid from each well and discard, making sure not to draw up the root. Add fresh 1% acetic acid and repeat this once or twice more to fully rinse stain from the root.
IV. Once rinsed, submerge each root in 1% acetic acid one final time, and leave to de-stain either for several hours or overnight.
V. Once the roots have destained in acetic acid, replace this with 50% glycerol and leave for a further couple of hours. This will leach out further stain from the root before mounting.

Mounting

I. Label each frosted slide at the top with the sample codes or names and the date. Each slide can hold two mounted root fragments measuring 1 cm in length.
II. For the first root, use a 200µl pipette to add 20-70ul 50% glycerol in a circle to the upper half of the slide. The amount of glycerol depends on the size and thickness of the root being mounted.
III. Using the dissection needle, carefully lift the root from the 12-well dish and place it into the glycerol. Gently use the staining needle to position the root so that no branches overlap.
IV. Slowly lower the coverslip over the glycerol, pinning the root in place. If there are air bubbles around the edges of the coverslip, use the 200µl pipette to carefully add enough glycerol to fill the space.
V. Once the coverslip is in position, gently dab away excess glycerol from the slide and add a dot of clear nail polish at each corner of the coverslip. Leave the coverslip to dry for at least ten minutes – during which the other roots can be mounted – and then seal the edges of the coverslip using more clear nail polish.
VI. Once sealed and dried, place the slides into a slide book.

Recipes for reagents

10% KOH

For 500ml of 10% KOH, dissolve 50g of KOH pellets in 500ml of dH₂O. Place a stir bar (flea) in the bottle and leave the solution on the stir plate until all the pellets have dissolved.

Staining solution (25% glacial acetic acid and 10% ink)

For 250ml of staining solution, prepare a stock solution of 25% glacial acetic acid by adding 125ml of glacial acetic acid to 500ml of dH₂O.

Then add 25ml of ink (either Sheaffer blue catalogue number: 94221 or Parker blue) to 225ml of the 25% acetic acid to produce 250ml of the staining solution. Mix well by inverting.

50% glycerol

For 50ml of 50% glycerol, mix 25ml of the stock glycerol solution with 25ml of dH₂O in a 50ml falcon tube. Mix the solution well by inverting.

2% HCl

Concentrated HCL should be around 38% and cannot exceed 40%. For 500ml of a 2% HCl solution, in a fume hood take 26.32ml of the concentrated HCl stock and dilute with 473.68ml of dH20.

Staining Woody Roots

- Try not to increase the time spent at 100, but roots can be left in 60°C water bath for much longer- for a couple of hours or more. The temperature can also be increased to 70°C but roots must be checked often.
- You can also leave roots in room temperature KOH for around 3 days. Check the colour of the solution regularly and refresh as needed
- During the staining step, roots may be left in the 100°C water bath in staining solution for less time; around 1 minute as opposed to 3 minutes.

**Methods S2****.** Image augmentation, balancing, calibration and contextualisation and model training set up.

*Image augmentation and balancing*

After individual images were split into training and test datasets and processed into 252x252 pixel (Arbuscular mycorrhizas, AM) or 126x126 pixel (Ericoid mycorrhizas, ErM) image tiles, the training datasets were augmented by randomly applying horizontal and vertical flips as well as changes in brightness and saturation (both by applying a multiplier randomly chosen from the interval [0.75, 1.25] to the relevant quantity). The image tiles were then randomly cropped with a probability of 30% to 90% of their original size (with a ratio of 1.0 per side) and subsequently resized to their original size to augment for a magnification-like effect (noting that this only approximates the effect of true magnification). Resulting augmented tiles were added to the original tiles to create a mixed training dataset consisting of both original and augmented tiles.

In addition to augmentation itself, additional measures were introduced within the image tile processing steps to counteract the imbalance across classes in the respective datasets. Depending on the class, an individual image tile was augmented into multiple tiles to increase the total number of tiles per given class. Table S2 lists the factors applied for both datasets. These factors reflect the number of times a tile was augmented and added to the total dataset exclusively, i.e. if a factor of 2 was applied to AM^+^, two augmented tiles were added along with the original image tile, resulting in three total tiles being added to the dataset. Second, an anchor class was defined for the respective datasets (AM^+^ for the AM dataset, Bl+ for the ErM dataset). The number of tiles for the anchor class was used as reference for the other corresponding classes as follows:

i) The number of tiles for the anchor class was multiplied by the appropriate “balance factor” (usually between 0.75 and 1.25; Table S2), resulting in the “anchor class number”.
ii) The number of tiles in each other class was reduced to the anchor class number by randomly deleting image tiles.
iii) Classes with a smaller number of tiles than the anchor class number were not altered.

This process enabled a significant balancing of the entire dataset with respect to the anchor class, as in most cases, heavily overrepresented classes such as N- or X were reduced. Note that the “balance factor” was also applied as a hyperparameter during hyperparameter sweeps (Tables S2, S3).

The resulting dataset was subsequently split into a training and validation set using an 80%/20% ratio. Unlike the division between test images and training/validation images, which applied on the whole-image basis, the subdivision between training and validation data was applied on an image tile basis. Furthermore, the splitting of training and validation sets was applied each time a training run was started, whereas the splitting of training and test data was static across all applied training runs.

### *Calibration*

As trained models might be overconfident or underconfident when classifying individual image tiles, model calibration following a “temperature scaling” approach (Guo et al. 2017) was applied to the best performing models. This resulted in a “temperature” scaling factor that was used to balance the output probabilities of a particular model to match its confidence on a prediction. To calibrate, the entire available datasets for AM and ErM, both training and test data, was used to yield the highest possible sample representation for each class. For exact values of the temperature factor, please refer to the source code.

Once each model was calibrated, the estimation of metrics and uncertainty across an image was based on Monte Carlo sampling per image tile. In this step, we consider the entire probability distribution provided by the calibrated model for each tile. By using repetitive sampling of these distributions across all tiles, a distribution of estimates for each metric within the image was generated. This approach aided the identification of the variability in predictions.

Finally, these sampling estimates have been used to calculate the mean and standard deviation to derive a confidence interval via:

- {Metric name} LowerConfidence = Mean – 2 * standard deviation
- {Metric name} UpperConfidence = Mean + 2 * standard deviation

This provides a quantitative indication of the relative confidence with which the model is able to predict a given metric for a given image, to be compared across the range of colonisation metrics and between images.

*Contextual Function*

Neighbouring tiles are of the same fixed size as the original image tile (252x252 pixels for AM and 126x126 for ErM) and each tile is run through the model separately, assigning a class prediction to all four overlapping image tiles (Fig. S3a). To achieve the highest potential improvement of the classification performance of the model, several degrees of diagonal overlap were tested using a *Macro F_1_* Score. Overlap values of 50%, 75% and 100% were tested, and an overlap of 75% was found to produce the best results. Corresponding model performances are listed in the Results section in the main manuscript.

*Model training set up*

All model training and testing was run on Microsoft’s Azure Machine Learning Studio platform. Model training was conducted using hyperparameter sweeps, with which several pre-defined hyperparameters (see below) were randomly initialised within a given interval. Each training trial was evaluated after having concluded 10 epochs for its validation loss and was subsequently reevaluated every five epochs. Trials that did not achieve a lower validation loss than the best trial in the given hyperparameter sweep were truncated using ratios between 33% - 50% to reduce required model optimisation time.

Hyperparameter sweeps were conducted on several candidate model architectures, including CNN1 (Evangelisti *et al.* 2021), ResNet50 (He *et al.* 2015), ResNeXt32x4 (Xie *et al.* 2015), EfficientNetB5 (Tan & Le 2019) and EfficientNetV2M (Tan & Le 2021). In addition to the balancing measures applied at augmentation stage, class weights following a distribution by effective number (Cui et al. 2019) were introduced in the loss function. Table S3 lists relevant parameters for all training runs conducted, whereby all runs applied a balanced loss function (with effective number class weights), as well as the respective augmentation factors. All hyperparameters followed a uniform distribution in the given intervals, except the batch size, for which a value was randomly chosen from the numbers given in the respective interval. Deeper model architectures, such as ResNet50 (He *et al.* 2015), ResNeXt32x4 (Xie *et al.* 2015) and EfficientNetB5 (Tan & Le 2019) were initialised in a pre-trained state (pre-trained on the ImageNet dataset, using the PyTorch configuration “IMAGENET1K_V1”). Thus, a lower training rate was applied compared to CNN1, which was initialised from scratch.

Figures S8 and S9 show trials applied for the EfficientNetB5 (Tan & Le 2019) architecture, and Table S4 summarises the hyperparameter intervals applied during hyperparameter sweep training on the AM and ErM dataset. As in the AM dataset, all hyperparameters followed a uniform distribution in the given interval, except the batch size where a random choice of the available value was applied instead. Similar pre-trained states as in the AM dataset were used for the EfficientNetB5 and the EfficientNetV2M architectures respectively. Hyperparameters such as balance factor, batch size and learning rate were slightly adjusted for the AM dataset, as they result in more consistent results and improved executions of the hyperparameter sweeps.

**Supplementary Tables**

**Table S1.** Best *Macro F_1_* Scores across all EfficientNet and EfficentNetV2 model architectures alongside best CNN1 models as a benchmark comparison for the arbuscular mycorrhiza (AM) and ericoid mycorrhiza (ErM) datasets. Note that the reported EfficientNetV2M model was trained on the same dataset, with added pure black and white images to the background class to make the model more robust against scanning artifacts.

| **Dataset** | **Model Architecture** | **Macro F_1_** |
| --- | --- | --- |
| AM | CNN1 – without contextual function | 47.54 |
|  | CNN1 – with contextual function | n/a |
|  | EfficientNetB5 – without contextual function | 63.32 |
|  | EfficientNetB5 – with contextual function | 66.47 |
|  | EfficientNetV2M – without contextual function | 60.30 |
|  | EfficientNetV2M – with contextual function | 64.57 |
| ErM | CNN1 – without contextual function | 40.61 |
|  | CNN1 – with contextual function | 42.94 |
|  | EfficientNetB5 – without contextual function | 53.89 |
|  | EfficientNetB5 – with contextual function | 57.03 |
|  | EfficientNetV2M – without contextual function | 55.50 |
|  | EfficientNetV2M – with contextual function | 57.39 |

**Table S2.** Augmentation factors applied to the individual tile classes during image tile augmentation. AM, arbuscular mycorrhiza; ErM, ericoid mycorrhiza. Tile classes explained in Table 2, NR corresponds to non-root background.

|  | AM: {AM^+^: 2, N-: 1, NR: 0, U: 3, DSE: 3, H: 3} |
| --- | --- |
|  | ErM: {Bl+: 2, Br+: 2, T+: 2, N-: 1, X: 0, M: 0, U: 3, D: 3, HE+: 3, HD+: 3} |

**Table S3.** Value intervals for hyperparameters considered in arbuscular mycorrhiza pathway of the model training via hyperparameter sweeps.

| Architecture | Pre-trained | Balance Factor | Batch Size | Learning Rate | Adam $\boldsymbol{\beta}_{\boldsymbol{1}}$ | Adam $\boldsymbol{\beta}_{\boldsymbol{2}}$ |
| --- | --- | --- | --- | --- | --- | --- |
| CNN1 | False | [0.75; 1.25] | [32, 64,  128, 256] | [1e-6; 1e-5] | [0.875; 0.925] | [0.950; 0.999] |
| ResNet50 | True | [0.75; 1.25] | [32, 64] | [5e-7; 5e-6] | [0.875; 0.925] | [0.950; 0.999] |
| ResNeXt32x4 | True | [0.75; 1.25] | [32, 64] | [5e-7; 5e-6] | [0.875; 0.925] | [0.950; 0.999] |
| EfficientNetB5 | True | [0.75; 1.25] | 32 | [5e-7; 5e-6] | [0.875; 0.925] | [0.950; 0.999] |

**Table S4:** Value intervals for hyperparameters considered in ericoid mycorrhizal pathway of the model training via hyperparameter sweeps.

| Architecture | Pre-trained | Balance Factor | Batch Size | Learning Rate | Adam $\boldsymbol{\beta}_{\boldsymbol{1}}$ | Adam $\boldsymbol{\beta}_{\boldsymbol{2}}$ |
| --- | --- | --- | --- | --- | --- | --- |
| CNN1 | False | [0.80; 1.20] | [32, 64] | [5e-6; 5e-5] | [0.875; 0.925] | [0.950; 0.999] |
| EfficientNetB5 | True | [0.80; 1.20] | 32 | [1e-6; 1e-5] | [0.875; 0.925] | [0.950; 0.999] |
| EfficientNetV2M | True | [0.80; 1.20] | 32 | [1e-6; 1e-5] | [0.875; 0.925] | [0.950; 0.999] |

### **Table S5.** Class determinations after running an ericoid field root sample (TW_05_Cal_01_04) through both the arbuscular and ericoid mycorrhiza pathways. Numbers indicate class tile counts followed by percent of root colonised by said class. 'na’, classes not including in the AM pathway.

| **Tile class** | **ErM path** | **AM path (determination/tile count)** | |
| --- | --- | --- | --- |
| Blue coils | 288 | AM^+^ | 25 |
| Brown coils | 236 |  | na |
| Type 2 | 0 |  | na |
| Mixed ErM | 7 | AM+DSE | 44 |
| ErM-DSE | 1 |  | na |
| Uncolonised | 666 |  | 601 |
| DSE | 17 |  | 565 |
| Mainroot | 2109 |  | na |
| Background | 62,206 |  | 14,979 |
| Colonised | 43.69% |  | 5.59% |
| Uncolonised | 54.91% |  | 48.66% |
| DSE | 1.40% |  | 45.75% |

**Table S6.** Training data for the arbuscular mycorrhiza (AM) model (MycorrhizaFinder). Numbers shown are the number of annotated tiles in each class.

| **Image name** | **AM^+^** | **N-** | **X** | **U** | **D** | **H** | **Total** |
| --- | --- | --- | --- | --- | --- | --- | --- |
| 10E_2L_E | 52 | 165 | 1163 | 0 | 0 | 0 | 1380 |
| 10E_2U_H | 68 | 57 | 946 | 11 | 25 | 43 | 1150 |
| 1B-3U-5 | 11 | 56 | 1380 | 28 | 17 | 8 | 1500 |
| 1B-3U-6 | 20 | 42 | 1138 | 150 | 23 | 22 | 1395 |
| 1D-1L-8 | 13 | 143 | 1748 | 61 | 57 | 2 | 2024 |
| 20C-2L-7 | 15 | 162 | 1838 | 12 | 62 | 2 | 2091 |
| 5C_3U_A | 102 | 139 | 1273 | 26 | 0 | 0 | 1540 |
| ACH700_C3_J-K_4 | 80 | 604 | 3408 | 8 | 48 | 0 | 4148 |
| ACH700_C2_O-P_15 | 58 | 108 | 4038 | 6 | 5 | 1 | 4216 |
| AFF756_C1_J-L_2 | 84 | 435 | 11444 | 14 | 7 | 0 | 11984 |
| high_res_white_image_AM | 0 | 0 | 4225 | 0 | 0 | 0 | 4225 |
| high_res_black_image_AM | 0 | 0 | 4225 | 0 | 0 | 0 | 4225 |
| AFF756_C1_J-L_3 | 86 | 412 | 11194 | 58 | 153 | 81 | 11984 |
| AFF756_C1_D-F_15 | 41 | 155 | 8908 | 3 | 5 | 14 | 9126 |
| ABS710_C3_J-L_6 | 44 | 310 | 3712 | 6 | 18 | 2 | 4092 |
| ZY820_C1_E-F_5_CROPPED | 201 | 640 | 115 | 122 | 11 | 3 | 1092 |
| ABS710_C1_E_2 | 236 | 1073 | 136 | 56 | 27 | 3 | 1531 |
| ABM841_C1_D_1 | 37 | 768 | 272 | 329 | 0 | 0 | 1406 |
| ABE784_C3_h_3 | 275 | 922 | 316 | 46 | 36 | 8 | 1603 |
| ACH700_C2_O-P_14 | 122 | 366 | 256 | 29 | 6 | 0 | 779 |
| ACH700_C2_A-B_15 | 101 | 620 | 11 | 217 | 1 | 0 | 950 |
| ABS710_C3_J-L_7 | 177 | 174 | 222 | 89 | 15 | 25 | 702 |
| ABM841_C2_A-B_4 | 109 | 622 | 26 | 1 | 20 | 2 | 780 |
| ABS710_C1_E_5 | 113 | 754 | 263 | 45 | 31 | 12 | 1218 |
| ABS710_C2_H_7 | 262 | 1000 | 1 | 24 | 16 | 2 | 1305 |
| ACH700_C2_A-B_4 | 75 | 609 | 108 | 46 | 18 | 0 | 856 |
| ACH700_C3_J-K_5 | 81 | 279 | 228 | 69 | 3 | 3 | 663 |
| ABM841_C2_A-B_5 | 86 | 566 | 60 | 12 | 8 | 0 | 732 |
| ACH700_C2_D_9 | 129 | 244 | 60 | 119 | 0 | 0 | 552 |
| ABS710_C3_J-L_2 | 43 | 194 | 159 | 0 | 20 | 2 | 418 |
| AFF756_C1_A-C_15 | 176 | 199 | 129 | 0 | 0 | 0 | 504 |
| ABE784_C3_h_4 | 151 | 391 | 38 | 38 | 4 | 1 | 623 |
| ABE784_C3_h_1 | 70 | 413 | 109 | 12 | 4 | 1 | 609 |
| ABM841_C2_A-B_10 | 36 | 242 | 15 | 29 | 0 | 0 | 322 |
| ADG811_C3_b_4 | 37 | 434 | 48 | 8 | 29 | 6 | 562 |
| ABM841_C1_D_2 | 8 | 340 | 31 | 0 | 44 | 0 | 423 |
| ABS710_C3_J-L_1 | 33 | 130 | 79 | 0 | 0 | 0 | 242 |
| ABS710_C3_P-Q_1 | 98 | 218 | 138 | 2 | 29 | 19 | 504 |
| ABS710_C3_P-Q_2 | 64 | 239 | 139 | 1 | 10 | 0 | 453 |
| ABS710_C3_J-L_3 | 15 | 55 | 64 | 4 | 40 | 11 | 189 |
| ABM841_C2_A-B_9 | 11 | 164 | 13 | 0 | 2 | 0 | 190 |
| AFF756_C1_D-F_16 | 2 | 364 | 0 | 5 | 0 | 0 | 371 |
| AFF756_C1_J-L_4 | 35 | 250 | 2 | 11 | 9 | 0 | 307 |
| ADG811-C2-K_1 | 107 | 129 | 11 | 4 | 5 | 3 | 259 |
| ABM841_C2_A-B_2 | 18 | 16 | 35 | 0 | 0 | 0 | 69 |
| ABS710_C3_J-L_10 | 5 | 25 | 7 | 4 | 0 | 0 | 41 |
| ABS710_C3_J-L_16 | 17 | 11 | 5 | 0 | 0 | 0 | 33 |

**Table S7.** Test data for the arbuscular mycorrhiza (AM) model (MycorrhizaFinder). Numbers shown are the number of annotated tiles in each class.

| **Image name** | **AM^+^** | **N-** | **X** | **U** | **D** | **H** | **Total** |
| --- | --- | --- | --- | --- | --- | --- | --- |
| 49B-2U-7 | 1 | 62 | 924 | 3 | 74 | 0 | 1064 |
| AFF756_C1_J-L_1 | 97 | 942 | 10621 | 119 | 114 | 91 | 11984 |
| ABS710_C3_J-L_4 | 101 | 159 | 3738 | 7 | 61 | 26 | 4092 |
| ZY820_C1_E-F_7 | 139 | 486 | 8159 | 168 | 16 | 0 | 8968 |
| 67D-2U-6 | 0 | 159 | 171 | 45 | 19 | 0 | 394 |
| ABS710_C2_H_4 | 200 | 750 | 243 | 100 | 10 | 1 | 1304 |
| ABS710_C2_H_6 | 254 | 316 | 3 | 36 | 3 | 9 | 621 |
| ABM841_C1_R-S_7 | 141 | 341 | 229 | 34 | 1 | 2 | 748 |
| ABS710_C3_J-L_12 | 81 | 196 | 22 | 29 | 0 | 0 | 328 |
| ABS710_C3_J-L_13 | 51 | 58 | 18 | 0 | 0 | 0 | 127 |

**Table S8.** Training data for the ericoid mycorrhiza (ErM) model (MycorrhizaFinder) . Numbers shown are the number of annotated tiles in each class.

| **Image name** | **Bl+** | **Br+** | **T+** | **N-** | **X** | **M** | **U** | **D** | **HE+** | **HD+** | **Total** |
| --- | --- | --- | --- | --- | --- | --- | --- | --- | --- | --- | --- |
| high_res_black_image_ErM | 0 | 0 | 0 | 0 | 16900 | 0 | 0 | 0 | 0 | 0 | 16900 |
| high_res_white_image_ErM | 0 | 0 | 0 | 0 | 16900 | 0 | 0 | 0 | 0 | 0 | 16900 |
| NO4_01_Cal_05-07_NO4_01_Cal_05 | 28 | 0 | 0 | 101 | 104 | 129 | 38 | 27 | 0 | 6 | 433 |
| NO4_01_Tet_01-04_NO4_01_Tet_04 | 17 | 8 | 0 | 94 | 1977 | 230 | 29 | 0 | 0 | 0 | 2355 |
| T1_14_2_01-04_T1_14_2_02 | 14 | 37 | 0 | 59 | 2042 | 314 | 147 | 0 | 2 | 0 | 2615 |
| NO2_09_Tet_05-08_NO2_09_Tet_07 | 30 | 13 | 0 | 44 | 129 | 77 | 94 | 0 | 5 | 0 | 392 |
| T1_16_3_01-04_T1_16_3_01 | 0 | 13 | 0 | 147 | 172 | 138 | 151 | 0 | 0 | 0 | 621 |
| T1_18_1_05-09_T1_18_1_06 | 101 | 12 | 0 | 102 | 105 | 157 | 156 | 0 | 2 | 1 | 636 |
| NO4_01_Tet_01-04_NO4_01_Tet_03 | 24 | 21 | 0 | 117 | 290 | 165 | 48 | 0 | 1 | 0 | 666 |
| T1_15_3_01-04_T1_15_3_01 | 61 | 21 | 0 | 95 | 162 | 349 | 16 | 0 | 4 | 0 | 708 |
| TW_06_Cal_01-04_TW_06_Cal_04 | 25 | 0 | 0 | 36 | 2385 | 27 | 12 | 0 | 0 | 0 | 2485 |
| NO1_1_Cal_1-4_NO1_1_Cal_4 | 0 | 183 | 26 | 28 | 261 | 536 | 24 | 0 | 1 | 0 | 1059 |
| NO2_09_Tet_05-08_NO2_09_Tet_06 | 33 | 5 | 0 | 17 | 71 | 95 | 164 | 0 | 2 | 0 | 387 |
| T1_17_1_01-04_T1_17_1_01 | 13 | 0 | 0 | 87 | 194 | 236 | 41 | 0 | 2 | 0 | 573 |
| T1_17_1_01-04_T1_17_1_03 | 13 | 65 | 0 | 39 | 150 | 179 | 126 | 0 | 8 | 0 | 580 |
| TW_04_Cal_01-04_TW_04_Cal_01 | 102 | 5 | 0 | 78 | 124 | 167 | 54 | 0 | 1 | 0 | 531 |
| T1_13_3_01-04_T1_13_3_01 | 0 | 71 | 0 | 58 | 149 | 154 | 14 | 0 | 0 | 0 | 446 |
| T1_13_3_01-04_T1_13_3_04 | 2 | 103 | 0 | 32 | 39 | 140 | 1 | 0 | 2 | 0 | 319 |
| NO1_1_Cal_1-4_NO1_1_Cal_3 | 2 | 101 | 0 | 146 | 112 | 278 | 1 | 0 | 2 | 0 | 642 |
| NO4_01_Tet_01-04_NO4_01_Tet_02 | 27 | 15 | 0 | 136 | 157 | 188 | 51 | 0 | 0 | 2 | 576 |
| NO1_09_Cal_01-04_NO1_09_Cal_02 | 35 | 0 | 0 | 159 | 137 | 133 | 33 | 0 | 0 | 0 | 497 |
| T1_13_1_01-04_T1_13_1_02 | 33 | 64 | 0 | 54 | 116 | 122 | 161 | 0 | 22 | 0 | 572 |
| NO1_06_Cal_01-04_NO1_06_Cal_02 | 48 | 9 | 0 | 23 | 1175 | 77 | 49 | 0 | 1 | 0 | 1382 |
| T1_16_2_01-04_T1_16_2_04 | 0 | 45 | 0 | 68 | 146 | 177 | 175 | 0 | 0 | 0 | 611 |
| NO4_01_Cal_05-07_NO4_01_Cal_06 | 46 | 3 | 0 | 47 | 87 | 25 | 23 | 37 | 0 | 0 | 268 |
| TW_01_Cal_01-04_TW_01_Cal_01 | 21 | 36 | 0 | 46 | 141 | 114 | 62 | 0 | 18 | 0 | 438 |
| T1_17_1_01-04_T1_17_1_02 | 5 | 38 | 0 | 90 | 151 | 222 | 36 | 0 | 1 | 0 | 543 |
| TW_05_Cal_05-07_TW_05_Cal_06 | 9 | 79 | 0 | 56 | 29 | 136 | 40 | 0 | 7 | 0 | 356 |
| NO1_04_Cal_01-04_NO1_4_Cal_1 | 14 | 1 | 94 | 151 | 115 | 126 | 6 | 0 | 0 | 0 | 507 |
| T1_16_1_01-04_T1_16_1_03 | 19 | 20 | 0 | 117 | 171 | 130 | 242 | 0 | 1 | 0 | 700 |
| T1_18_3_01-04_T1_18_3_02 | 8 | 59 | 0 | 85 | 133 | 125 | 120 | 0 | 3 | 0 | 533 |
| T1_13_1_01-04_T1_13_1_04 | 30 | 11 | 0 | 76 | 44 | 152 | 45 | 0 | 3 | 0 | 361 |
| TW_06_Cal_01-04_TW_06_Cal_03 | 63 | 0 | 0 | 130 | 96 | 370 | 2 | 0 | 0 | 0 | 661 |
| T1_16_2_01-04_T1_16_2_03 | 0 | 47 | 0 | 88 | 124 | 127 | 267 | 0 | 0 | 0 | 653 |
| AM_1D-1L-8 | 0 | 0 | 0 | 0 | 0 | 0 | 0 | 102 | 0 | 0 | 102 |
| TW_02_Cal_01-04_TW_02_Cal_02 | 24 | 49 | 0 | 159 | 199 | 293 | 149 | 0 | 4 | 0 | 877 |
| NO2_09_Tet_05-08_NO2_09_Tet_05 | 62 | 32 | 0 | 117 | 172 | 102 | 164 | 0 | 9 | 0 | 658 |
| NO4_01_Cal_05-07_NO4_01_Cal_07 | 0 | 0 | 0 | 0 | 71 | 0 | 0 | 78 | 0 | 14 | 163 |
| NO4_01_Tet_01-04_NO4_01_Tet_01 | 0 | 24 | 0 | 211 | 159 | 244 | 15 | 0 | 0 | 0 | 653 |
| T1_16_3_01-04_T1_16_3_04 | 0 | 0 | 0 | 296 | 170 | 188 | 88 | 0 | 0 | 0 | 742 |
| T1_14_2_01-04_T1_14_2_04 | 7 | 30 | 0 | 72 | 53 | 99 | 51 | 0 | 0 | 0 | 312 |
| T1_17_1_01-04_T1_17_1_04 | 10 | 24 | 0 | 34 | 110 | 142 | 182 | 0 | 0 | 0 | 502 |
| NO1_02_Tet_01-04_NO1_02_Tet_02 | 68 | 11 | 0 | 70 | 45 | 140 | 24 | 0 | 4 | 0 | 362 |
| TW_06_Cal_01-04_TW_06_Cal_02 | 15 | 3 | 0 | 158 | 69 | 205 | 0 | 0 | 0 | 0 | 450 |
| T1_18_1_05-09_T1_18_1_07 | 6 | 21 | 0 | 35 | 64 | 88 | 34 | 0 | 0 | 0 | 248 |
| TW_02_Cal_05-08_TW_02_Cal_06 | 51 | 18 | 0 | 18 | 43 | 91 | 34 | 0 | 12 | 0 | 267 |
| NO1_09_Tet_01-04_NO1_09_Tet_04 | 35 | 27 | 0 | 104 | 104 | 130 | 37 | 0 | 16 | 0 | 453 |
| NO4_01_Cal_01-04_NO4_01_Cal_02 | 43 | 3 | 9 | 51 | 89 | 43 | 39 | 3 | 2 | 2 | 284 |
| T1_15_3_01-04_T1_15_3_03 | 15 | 25 | 0 | 123 | 151 | 147 | 89 | 0 | 0 | 0 | 550 |
| T1_13_1_01-04_T1_13_1_01 | 46 | 51 | 0 | 51 | 138 | 116 | 31 | 0 | 12 | 0 | 445 |
| NO1_09_Cal_01-04_NO1_09_Cal_03 | 151 | 0 | 0 | 76 | 217 | 207 | 23 | 0 | 0 | 0 | 674 |
| NO1_09_Cal_01-04_NO1_09_Cal_04 | 21 | 15 | 0 | 68 | 87 | 85 | 65 | 0 | 0 | 0 | 341 |
| NO2_05_Tet_05-08_NO2_05_Tet_05 | 61 | 15 | 0 | 47 | 101 | 155 | 52 | 0 | 19 | 0 | 450 |
| TW_03_Cal_01-04_TW_03_Cal_04 | 28 | 51 | 0 | 44 | 71 | 143 | 52 | 0 | 5 | 0 | 394 |
| AM_ABS710_C3_J-L_4 | 0 | 0 | 0 | 0 | 0 | 0 | 0 | 141 | 0 | 0 | 141 |
| T1_16_1_01-04_T1_16_1_01 | 18 | 29 | 0 | 159 | 119 | 188 | 172 | 0 | 3 | 0 | 688 |
| AM_AFF756_C1_J-L_3 | 0 | 0 | 0 | 0 | 0 | 0 | 0 | 369 | 0 | 0 | 369 |
| T1_16_2_01-04_T1_16_2_02 | 0 | 94 | 0 | 46 | 59 | 146 | 57 | 0 | 0 | 0 | 402 |
| NO4_02_Tet_01-04_EnhancedColors_Extended_NO4_02_Tet_02 | 34 | 6 | 5 | 99 | 111 | 131 | 57 | 1 | 2 | 2 | 448 |
| TW_02_Cal_01-04_TW_02_Cal_04 | 12 | 54 | 0 | 57 | 128 | 79 | 14 | 0 | 4 | 0 | 348 |
| TW_04_Cal_01-04_TW_04_Cal_03 | 58 | 12 | 0 | 79 | 132 | 184 | 30 | 0 | 6 | 0 | 501 |
| NO1_04_Cal_01-04_NO1_4_Cal_3 | 29 | 8 | 1 | 106 | 129 | 132 | 0 | 0 | 0 | 0 | 405 |
| T1_13_1_01-04_T1_13_1_03 | 21 | 26 | 0 | 41 | 55 | 118 | 93 | 0 | 3 | 0 | 357 |
| NO1_09_Tet_01-04_NO1_09_Tet_01 | 0 | 0 | 142 | 15 | 46 | 182 | 54 | 0 | 0 | 0 | 439 |
| NO1_02_Tet_01-04_NO1_02_Tet_03 | 54 | 7 | 0 | 42 | 109 | 116 | 23 | 0 | 0 | 0 | 351 |
| NO1_02_Tet_01-04_NO1_02_Tet_01 | 23 | 2 | 1 | 76 | 84 | 112 | 16 | 0 | 2 | 0 | 316 |
| NO2_05_Tet_05-08_NO2_05_Tet_06 | 3 | 80 | 0 | 28 | 108 | 58 | 123 | 0 | 0 | 0 | 400 |
| T1_16_3_01-04_T1_16_3_03 | 45 | 36 | 0 | 67 | 112 | 56 | 64 | 0 | 2 | 0 | 382 |
| T1_18_3_01-04_T1_18_3_03 | 28 | 25 | 0 | 39 | 128 | 226 | 54 | 0 | 9 | 0 | 509 |
| T1_16_1_01-04_T1_16_1_02 | 40 | 28 | 0 | 71 | 127 | 162 | 153 | 0 | 6 | 0 | 587 |
| TW_05_Cal_01-04_EnhancedColors_Extended_TW_05_Cal_01 | 27 | 41 | 0 | 77 | 109 | 218 | 50 | 0 | 5 | 0 | 527 |
| TW_01_Cal_01-04_TW_01_Cal_04 | 38 | 22 | 0 | 52 | 89 | 123 | 40 | 1 | 7 | 0 | 372 |
| TW_03_Cal_01-04_TW_03_Cal_03 | 3 | 57 | 0 | 60 | 69 | 119 | 69 | 0 | 5 | 0 | 382 |
| T1_18_3_01-04_T1_18_3_04 | 16 | 15 | 0 | 60 | 83 | 240 | 47 | 0 | 2 | 0 | 463 |
| TW_05_Cal_01-04_EnhancedColors_Extended_TW_05_Cal_02 | 36 | 26 | 0 | 45 | 97 | 115 | 26 | 0 | 6 | 0 | 351 |
| NO1_06_Cal_01-04_NO1_06_Cal_04 | 8 | 46 | 0 | 62 | 43 | 126 | 44 | 0 | 2 | 0 | 331 |
| AM_1B-3U-5 | 0 | 0 | 0 | 0 | 0 | 0 | 0 | 36 | 0 | 0 | 36 |
| T1_16_2_01-04_T1_16_2_01 | 1 | 75 | 0 | 63 | 63 | 68 | 0 | 0 | 1 | 0 | 271 |
| AM_AFF756_C1_J-L_1 | 0 | 0 | 0 | 0 | 0 | 0 | 0 | 284 | 0 | 0 | 284 |
| T1_14_2_01-04_T1_14_2_03 | 4 | 31 | 0 | 78 | 72 | 163 | 165 | 0 | 2 | 0 | 515 |
| T1_14_2_05-08_T1_14_2_06 | 10 | 31 | 0 | 24 | 69 | 136 | 70 | 0 | 2 | 0 | 342 |
| NO1_04_Cal_01-04_NO1_4_Cal_4 | 37 | 9 | 0 | 57 | 91 | 104 | 0 | 0 | 8 | 0 | 306 |
| TW_01_Cal_01-04_TW_01_Cal_02 | 35 | 18 | 0 | 76 | 151 | 140 | 53 | 0 | 7 | 0 | 480 |
| NO1_06_Cal_01-04_NO1_06_Cal_01 | 15 | 42 | 0 | 42 | 101 | 118 | 30 | 0 | 9 | 0 | 357 |
| NO5_07_Cal_01-04_NO5_07_Cal_01 | 4 | 61 | 0 | 49 | 116 | 118 | 15 | 31 | 4 | 6 | 404 |
| TW_04_Cal_05-08_TW_04_Cal_05 | 75 | 3 | 0 | 77 | 167 | 153 | 30 | 0 | 0 | 0 | 505 |
| T1_15_3_01-04_T1_15_3_04 | 3 | 40 | 0 | 53 | 73 | 94 | 11 | 0 | 7 | 0 | 281 |
| AM_ABS710_C3_J-L_3 | 0 | 0 | 0 | 0 | 0 | 0 | 0 | 73 | 0 | 0 | 73 |
| AM_ACH700_C3_J-K_4 | 0 | 0 | 0 | 0 | 0 | 0 | 0 | 74 | 0 | 0 | 74 |
| NO1_06_Cal_01-04_NO1_06_Cal_03 | 31 | 12 | 20 | 14 | 106 | 118 | 20 | 0 | 29 | 0 | 350 |
| T1_13_3_01-04_T1_13_3_03 | 43 | 21 | 0 | 85 | 212 | 193 | 1 | 0 | 4 | 0 | 559 |
| T1_13_3_01-04_T1_13_3_02 | 49 | 11 | 0 | 41 | 58 | 95 | 7 | 0 | 5 | 0 | 266 |
| AM_67D-2U-6 | 0 | 0 | 0 | 0 | 0 | 0 | 0 | 25 | 0 | 0 | 25 |
| NO1_09_Tet_01-04_NO1_09_Tet_02 | 41 | 0 | 21 | 31 | 66 | 134 | 41 | 0 | 0 | 0 | 334 |
| TW_02_Cal_01-04_TW_02_Cal_03 | 16 | 11 | 0 | 81 | 76 | 159 | 56 | 0 | 1 | 0 | 400 |
| AM_ABM841_C1_D_2 | 0 | 0 | 0 | 0 | 0 | 0 | 0 | 84 | 0 | 0 | 84 |
| AM_ABS710_C3_J-L_7 | 0 | 0 | 0 | 0 | 0 | 0 | 0 | 37 | 0 | 0 | 37 |
| NO5_10_Cal_01-04_EnhancedColors_Extended_NO5_10_Cal_02 | 21 | 22 | 0 | 100 | 5 | 1 | 0 | 28 | 0 | 3 | 180 |
| AM_ABM841_C2_A-B_4 | 0 | 0 | 0 | 0 | 0 | 0 | 0 | 37 | 0 | 0 | 37 |
| AM_ABS710_C1_E_5 | 0 | 0 | 0 | 0 | 0 | 0 | 0 | 57 | 0 | 0 | 57 |
| AM_ABS710_C3_J-L_2 | 0 | 0 | 0 | 0 | 0 | 0 | 0 | 28 | 0 | 0 | 28 |
| AM_ABS710_C1_E_2 | 0 | 0 | 0 | 0 | 0 | 0 | 0 | 50 | 0 | 0 | 50 |
| AM_ABE784_C3_h_3 | 0 | 0 | 0 | 0 | 0 | 0 | 0 | 51 | 0 | 0 | 51 |
| AM_ABS710_C3_P-Q_1 | 0 | 0 | 0 | 0 | 0 | 0 | 0 | 50 | 0 | 0 | 50 |
| AM_ACH700_C2_A-B_4 | 0 | 0 | 0 | 0 | 0 | 0 | 0 | 28 | 0 | 0 | 28 |
| NO2_09_Tet_01-04_EnhancedColors_Extended_NO2_09_Tet_Frag | 0 | 0 | 0 | 0 | 0 | 0 | 0 | 4 | 0 | 0 | 4 |
| AM_ZY820_C1_E-F_5_CROPPED | 0 | 0 | 0 | 0 | 0 | 0 | 0 | 20 | 0 | 0 | 20 |
| AM_ACH700_C3_J-K_5 | 0 | 0 | 0 | 0 | 0 | 0 | 0 | 10 | 0 | 0 | 10 |
| AM_ACH700_C2_O-P_14 | 0 | 0 | 0 | 0 | 0 | 0 | 0 | 10 | 0 | 0 | 10 |
| AM_ABS710_C2_H_4 | 0 | 0 | 0 | 0 | 0 | 0 | 0 | 16 | 0 | 0 | 16 |
| AM_ABM841_C2_A-B_5 | 0 | 0 | 0 | 0 | 0 | 0 | 0 | 9 | 0 | 0 | 9 |
| AM_ACH700_C2_O-P_15 | 0 | 0 | 0 | 0 | 0 | 0 | 0 | 8 | 0 | 0 | 8 |
| AM_AFF756_C1_D-F_15 | 0 | 0 | 0 | 0 | 0 | 0 | 0 | 17 | 0 | 0 | 17 |
| AM_ABS710_C3_P-Q_2 | 0 | 0 | 0 | 0 | 0 | 0 | 0 | 15 | 0 | 0 | 15 |
| AM_ABS710_C2_H_6 | 0 | 0 | 0 | 0 | 0 | 0 | 0 | 8 | 0 | 0 | 8 |
| NO2_09_Tet_01-04_EnhancedColors_Extended_NO2_09_Tet_03 | 0 | 0 | 0 | 0 | 0 | 0 | 0 | 20 | 0 | 1 | 21 |
| AM_ADG811-C2-K_1 | 0 | 0 | 0 | 0 | 0 | 0 | 0 | 12 | 0 | 0 | 12 |
| AM_AFF756_C1_J-L_4 | 0 | 0 | 0 | 0 | 0 | 0 | 0 | 13 | 0 | 0 | 13 |
| AM_AFF756_C1_J-L_2 | 0 | 0 | 0 | 0 | 0 | 0 | 0 | 11 | 0 | 0 | 11 |
| AM_ABE784_C3_h_1 | 0 | 0 | 0 | 0 | 0 | 0 | 0 | 4 | 0 | 0 | 4 |
| AM_ABE784_C3_h_4 | 0 | 0 | 0 | 0 | 0 | 0 | 0 | 4 | 0 | 0 | 4 |
| AM_ABM841_C2_A-B_9 | 0 | 0 | 0 | 0 | 0 | 0 | 0 | 2 | 0 | 0 | 2 |
| AM_ACH700_C2_A-B_15 | 0 | 0 | 0 | 0 | 0 | 0 | 0 | 1 | 0 | 0 | 1 |

**Table S9.** Test data for the ericoid mycorrhiza (ErM) model (MycorrhizaFinder). Numbers shown are the number of annotated tiles in each class.

| **Image name** | **Bl+** | **Br+** | **T+** | **N-** | **X** | **M** | **U** | **D** | **HE+** | **HD+** | **Total** |
| --- | --- | --- | --- | --- | --- | --- | --- | --- | --- | --- | --- |
| T1_16_3_01-04_T1_16_3_02 | 30 | 84 | 0 | 81 | 2400 | 157 | 153 | 0 | 3 | 0 | 2908 |
| TW_04_Cal_01-04_TW_04_Cal_04 | 63 | 10 | 0 | 51 | 2442 | 126 | 19 | 0 | 6 | 0 | 2717 |
| TW_04_Cal_01-04_TW_04_Cal_02 | 24 | 0 | 0 | 63 | 2424 | 111 | 42 | 0 | 1 | 0 | 2665 |
| NO1_09_Cal_01-04_NO1_09_Cal_01 | 115 | 0 | 0 | 118 | 2300 | 218 | 46 | 0 | 0 | 0 | 2797 |
| AM_49B-2U-7 | 0 | 0 | 0 | 0 | 0 | 0 | 0 | 118 | 0 | 0 | 118 |
| NO4_01_Cal_01-04_NO4_01_Cal_01 | 27 | 0 | 0 | 74 | 84 | 101 | 64 | 20 | 0 | 2 | 372 |
| AM_10E_2U_H | 0 | 0 | 0 | 0 | 0 | 0 | 0 | 89 | 0 | 0 | 89 |
| T1_15_3_01-04_T1_15_3_02 | 3 | 23 | 0 | 131 | 161 | 145 | 43 | 0 | 0 | 0 | 506 |
| NO2_09_Tet_05-08_NO2_09_Tet_08 | 17 | 17 | 0 | 17 | 56 | 142 | 4 | 0 | 2 | 0 | 255 |
| TW_01_Cal_01-04_TW_01_Cal_03 | 29 | 23 | 0 | 52 | 92 | 137 | 90 | 0 | 5 | 0 | 428 |
| NO1_1_Cal_1-4_NO1_1_Cal_2 | 2 | 132 | 0 | 61 | 169 | 208 | 24 | 0 | 3 | 0 | 599 |
| AM_1B-3U-6 | 0 | 0 | 0 | 0 | 0 | 0 | 0 | 76 | 0 | 0 | 76 |
| AM_20C-2L-7 | 0 | 0 | 0 | 0 | 0 | 0 | 0 | 110 | 0 | 0 | 110 |
| NO1_04_Cal_01-04_NO1_4_Cal_2 | 10 | 29 | 60 | 182 | 215 | 228 | 54 | 0 | 0 | 0 | 778 |
| T1_14_2_05-08_T1_14_2_05 | 18 | 28 | 0 | 58 | 118 | 168 | 143 | 0 | 6 | 0 | 539 |
| T1_15_1_05-06_T1_15_1_06 | 19 | 28 | 0 | 72 | 53 | 130 | 132 | 0 | 4 | 0 | 438 |
| T1_15_1_05-06_T1_15_1_05 | 2 | 20 | 0 | 165 | 184 | 174 | 120 | 0 | 0 | 0 | 665 |
| TW_02_Cal_01-04_TW_02_Cal_01 | 89 | 11 | 0 | 80 | 106 | 205 | 98 | 0 | 3 | 0 | 593 |
| T1_18_3_01-04_T1_18_3_01 | 0 | 58 | 0 | 51 | 155 | 175 | 71 | 0 | 1 | 0 | 511 |
| TW_05_Cal_01-04_EnhancedColors_Extended_TW_05_Cal_04 | 16 | 0 | 0 | 47 | 81 | 79 | 0 | 0 | 0 | 0 | 223 |
| NO1_1_Cal_1-4_NO1_1_Cal_1 | 10 | 66 | 0 | 132 | 207 | 223 | 35 | 0 | 0 | 0 | 673 |
| NO4_01_Cal_01-04_NO4_01_Cal_03 | 53 | 0 | 0 | 35 | 73 | 77 | 56 | 6 | 0 | 7 | 307 |
| NO1_09_Tet_01-04_NO1_09_Tet_03 | 31 | 0 | 54 | 57 | 118 | 136 | 54 | 0 | 7 | 0 | 457 |
| TW_05_Cal_01-04_EnhancedColors_Extended_TW_05_Cal_03 | 39 | 6 | 0 | 38 | 138 | 89 | 41 | 0 | 3 | 0 | 354 |
| T1_16_1_01-04_T1_16_1_04 | 20 | 36 | 0 | 62 | 116 | 200 | 134 | 0 | 3 | 0 | 571 |
| TW_06_Cal_01-04_TW_06_Cal_01 | 20 | 0 | 0 | 417 | 1 | 0 | 35 | 0 | 0 | 0 | 473 |
| AM_ABS710_C3_J-L_6 | 0 | 0 | 0 | 0 | 0 | 0 | 0 | 32 | 0 | 0 | 32 |
| AM_ZY820_C1_E-F_7 | 0 | 0 | 0 | 0 | 0 | 0 | 0 | 33 | 0 | 0 | 33 |
| AM_ABS710_C2_H_7 | 0 | 0 | 0 | 0 | 0 | 0 | 0 | 27 | 0 | 0 | 27 |
| AM_ABM841_C1_R-S_7 | 0 | 0 | 0 | 0 | 0 | 0 | 0 | 1 | 0 | 0 | 1 |
